# Active Time-Restricted Feeding Protects the Blood Pressure Circadian Rhythm in Diabetic Mice

**DOI:** 10.1101/2020.07.17.209007

**Authors:** Tianfei Hou, Wen Su, Marilyn J. Duncan, Vsevolozhskaya A. Olga, Zhenheng Guo, Ming C. Gong

**Author notes:** To whom correspondence should be addressed: Ming C. Gong, Ph.D., 509 Wethington Building, 900 South Limestone Street, Lexington, KY 40536, Zhenheng Guo, Ph.D., 515 Wethington Building, 900 South Limestone Street, Lexington, KY 40536.

## Abstract

The quantity and quality of food intake have been considered crucial for peoples’ wellness. Only recently has it become appreciated that the timing of food intake, independent of total calorie consumption, is also critical. The involvement and contribution of the timing of food intake in nondipping blood pressure (BP), the most common disruption of the BP circadian rhythm in diabetes, however, remains uncertain. Here, we demonstrate that the loss of diurnal rhythm in food intake coincided with nondipping BP in type 2 diabetic *db/db* mice, and imposing a food intake diurnal rhythm by active time-restricted feeding (ATRF) remarkably prevented *db/db* mice from the development of nondipping BP. Moreover, ATRF, independent of calorie restriction, also effectively restored the already disrupted BP circadian rhythm in *db/db* mice. Mechanistically, ATRF reduced the sympathetic vascular tone and BP during the light phase via fasting, thus protecting the BP circadian rhythm in *db/db* mice. Moreover, we identified BMAL1, an obligatory core clock gene, as a potentially key molecule that links ATRF, sympathetic vascular tone, and BP circadian rhythm. Collectively, these data reveal an important but previously unrecognized role of the timing of food intake in the onset, prevention, and treatment of nondipping BP in diabetes.

## Introduction

As the incidence of type 2 diabetes mellitus continues to rise, diabetes has become one of the most prevalent and costly chronic diseases worldwide (1). About 75% of type 2 diabetic patients have hypertension (2), which worsens diabetic vascular complications and significantly contributes to their morbidity and mortality (3). With the increased use of ambulatory blood pressure (BP) monitoring, accumulating evidence indicates that not only the level but also the circadian rhythm of BP are critical for cardiovascular health (4). BP exhibits a circadian rhythm, with a rise during the early morning and a higher level maintained throughout the day, followed by a decrease of about 10%–20% at night (dip). Nondipping BP, defined as a less than 10% nocturnal decline in BP (4), is the most common disruption of the BP circadian rhythm in type 2 diabetes (5). The prevalence of nondipping BP varies among patient populations but is as high as 73% in type 2 diabetic patients (5). Nondipping BP, independent of hypertension, is associated with left ventricular hypertrophy, increased proteinuria, secondary forms of hypertension, increased insulin resistance, and increased fibrinogen level (6). Moreover, a recent large meta-analysis of 17,312 hypertensive individuals with 4 to 8 years follow-up revealed that nondippers, relative to dippers, have a significant 27% higher risk for total cardiovascular events (7). Despite the risk associated with nondipping BP in diabetic patients, the mechanisms underlying this problem are largely unknown. As a result, there is currently no effective medication for the prevention and treatment of nondipping BP in diabetes.

It is well known that excessive food intake is a major risk factor for the onset of obesity and type 2 diabetes (8). Research over the past decade has demonstrated that the quantity of food intake (how much we eat) and/or the quality of food intake (what we eat) has a profound impact on obesity and diabetes (8). Only recently has it become appreciated that the timing of food intake (when we eat), independent of total caloric and macronutrient quality in foods (9), is also critical for diet-induced metabolic diseases. Observational human studies revealed that late-night eating is higher in obese and diabetic patients compared to healthy individuals (10) and is associated with the development of metabolic disorders (11). In line with these human studies, animal work further demonstrated a causal relationship between the timing of food intake and the development of obesity and metabolic diseases. Of interest, when mice are fed an obesogenic diet *ad libitum* (ALF), they develop obesity and metabolic disorders (12, 13). In contrast, when mice are fed the same diet only during the active phase (active time-restricted feeding [ATRF]), they do not develop obesity and metabolic disorders (12, 13). Together, these studies suggest that the timing of food intake, independent of calorie restriction, has a profound impact on metabolic health. However, the relationship between the timing of food intake and BP circadian rhythm is unknown, and whether ATRF can serve as a therapeutic strategy against nondipping BP in diabetes is also unknown.

The current study seeks to investigate the cause and effect relationship between the timing of food intake and BP circadian rhythm in type 2 diabetic *db/db* mice. Our results show that the loss of food intake diurnal rhythm coincides with nondipping BP in *db/db* mice and that imposing a food intake diurnal rhythm by ATRF, independent of calorie restriction, effectively protects BP circadian rhythm in *db/db* mice. Mechanistically, we demonstrate that ATRF protects BP circadian rhythm via suppressing the sympathetic vascular tone during the fasting light phase in *db/db* mice and identify BMAL1, an obligatory core clock gene, as a key molecular player in the ATRF protection of BP circadian rhythm. Our results reveal an important but previously unrecognized role of the timing of food intake in BP circadian rhythm in diabetes.

## Results

### Loss of food intake diurnal rhythm coincides with nondipping BP in *db/db* mice

We and others have reported that the food intake diurnal rhythm (14) and BP circadian rhythm (15-17) are altered in type 2 diabetic *db/db* mice. To investigate whether the altered food intake diurnal rhythm is involved in nondipping BP in diabetes, we simultaneously monitored food intake and BP circadian rhythms by BioDAQ and telemetry in the same *db/db* mice and age-matched nondiabetic control (*db/+*) mice at 9-, 10-, 11-, and 12 weeks of age. BioDAQ is a computer-automated instrument that can be programmed to control the gates to food chambers at specified times and to monitor the amount and pattern of food intake continuously. As expected, the *db/db* mice consumed more food than the control mice throughout 9 to 12 weeks of age (Figure 1A). Consequently, the *db/db* mice developed obesity and hyperglycemia (Supplemental Figure 1A and 1B). Analysis of daily food intake revealed that the increase in food intake was more dramatic during the light phase than the dark phase in the *db/db* mice relative to that in the control mice (Figure 1B). As such, the diurnal rhythm of food intake was significantly dampened in the *db/db* mice throughout 9-12 weeks of age (Figure 1C).

**Figure 1.**
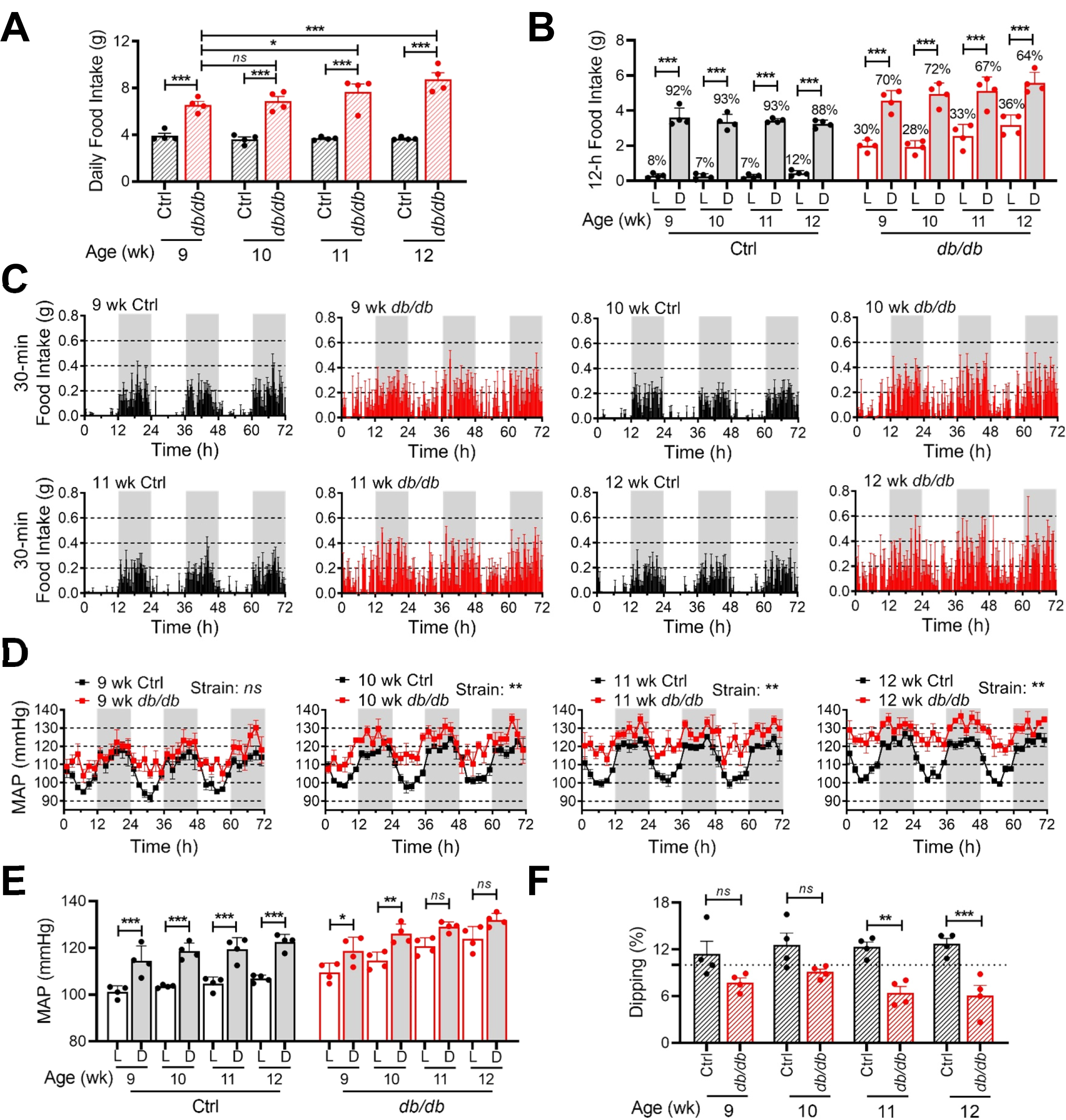
Time course of the food intake diurnal rhythm and blood pressure circadian rhythm in *db/db* and control mice. Food intake and blood pressure were recorded by BioDAQ and telemetry in the same 9-, 10-, 11-, and 12-week (wk)-old type 2 diabetic *db/db* mice and age-matched nondiabetic control mice (*db/+*). Four mice in each group. (**A**) Daily food intake. (**B**) The 12-hour (h) food intake during the light (L) and dark (D) phases. Percent of daily food intakes during the light and dark phase are indicated above each bar. (**C**) Daily profiles of food intake in 30-minute intervals over 72 hours (h) during the light and dark phases, shown in white and gray, respectively. (**D**) Daily profiles of mean arterial pressure (MAP) over 72 h in 2-h intervals during the light and dark phases. (**E**) The 12-h average MAP during the light (L) and dark (D) phases. (**F**) MAP dipping was calculated as % of MAP decrease during the light phase compared to the dark phase. The dashed line indicates 10% dipping. The data were analyzed by 2-way ANOVA (A, D, and F) and 3-way ANOVA (B and E) with multiple comparisons test. *, p<0.05; **, p<0.01; ***, p<0.001; ns, not significant.

The circadian rhythm in mean arterial pressure (MAP) was monitored by telemetry in these conscious free-moving *db/db* and control mice at the same time when their food intake was monitored. Consistent with our previous reports (15, 16), a robust MAP circadian rhythm was found in the control mice but gradually diminished in the *db/db* mice in an age-dependent manner (Figure 1D). Interestingly, the diminishment of BP circadian rhythm in the *db/db* mice primarily resulted from the larger amplitude of MAP during the light phase than during the dark phase (Figure 1D), which coincided with the increased food intake during the light phase in the *db/db* mice (Figure 1B and 1C). In the *db/db* mice, the daytime elevation of MAP was evident as early as 9 weeks of age before the nighttime elevation of MAP occurred (Figure 1D). As a result, the normal diurnal MAP variation, as seen in control mice, was blunted in the *db/db* mice in an age-dependent manner (Figure 1E). Moreover, the *db/db* mice developed nondipping BP by 9 weeks of age (Figure 1F).

### Imposing a food intake diurnal rhythm by ATRF prevents *db/db* mice from the development of nondipping BP

The concomitant attenuation of the food intake diurnal rhythm and BP circadian rhythm indicates that the attenuation in the food intake diurnal rhythm may cause nondipping BP in *db/db* mice. To test this possibility, we investigated whether imposing a food intake diurnal rhythm by ATRF prevents *db/db* mice from nondipping BP. The 6-week-old *db/db* and control mice were fed a chow diet *ad libitum* feeding (ALF) or ATRF (food was available only during the active dark phase for 8 hours [hrs]). Food intake and BP were measured by indirect calorimetry and telemetry, respectively, after 10 weeks of ATRF or ALF.

Representative telemetry recordings (Figure 2A) and quantitative data (Figure 2B) illustrate that the control mice under ALF exhibited a clear-cut MAP circadian rhythm, characterized by a higher MAP during the dark phase than during the light phase, which correlated with their food intake diurnal rhythm (Supplemental Figure 2A). In contrast, the *db/db* mice under ALF exhibited an impaired MAP circadian rhythm (Figure 2A and 2B), abolished diurnal MAP variation (Figure 2C), and nondipping MAP (Figure 2D), which correlated with the diminished food intake diurnal rhythm (Supplemental Figure 2B *vs.* 2A). Remarkably, imposing a food intake diurnal rhythm by ATRF (Supplemental 2D) prevented the *db/db* mice from the development of impaired MAP circadian rhythm, abolished diurnal MAP variation, and nondipping MAP, but did not significantly affect these parameters in the control mice (Supplemental 2C and Figure 2A–2D). Importantly, ATRF selectively dwindled the MAP during the light phase while the *db/db* mice were fasting, whereas ATRF had little effect on the MAP during the dark phase when the *db/db* mice were feeding (Figure 2A–2C; Supplemental Figure 2C and 2D). These data suggest that ATRF protects *db/db* mice from the development of nondipping BP by inducing fasting during the light phase.

**Figure 2.**
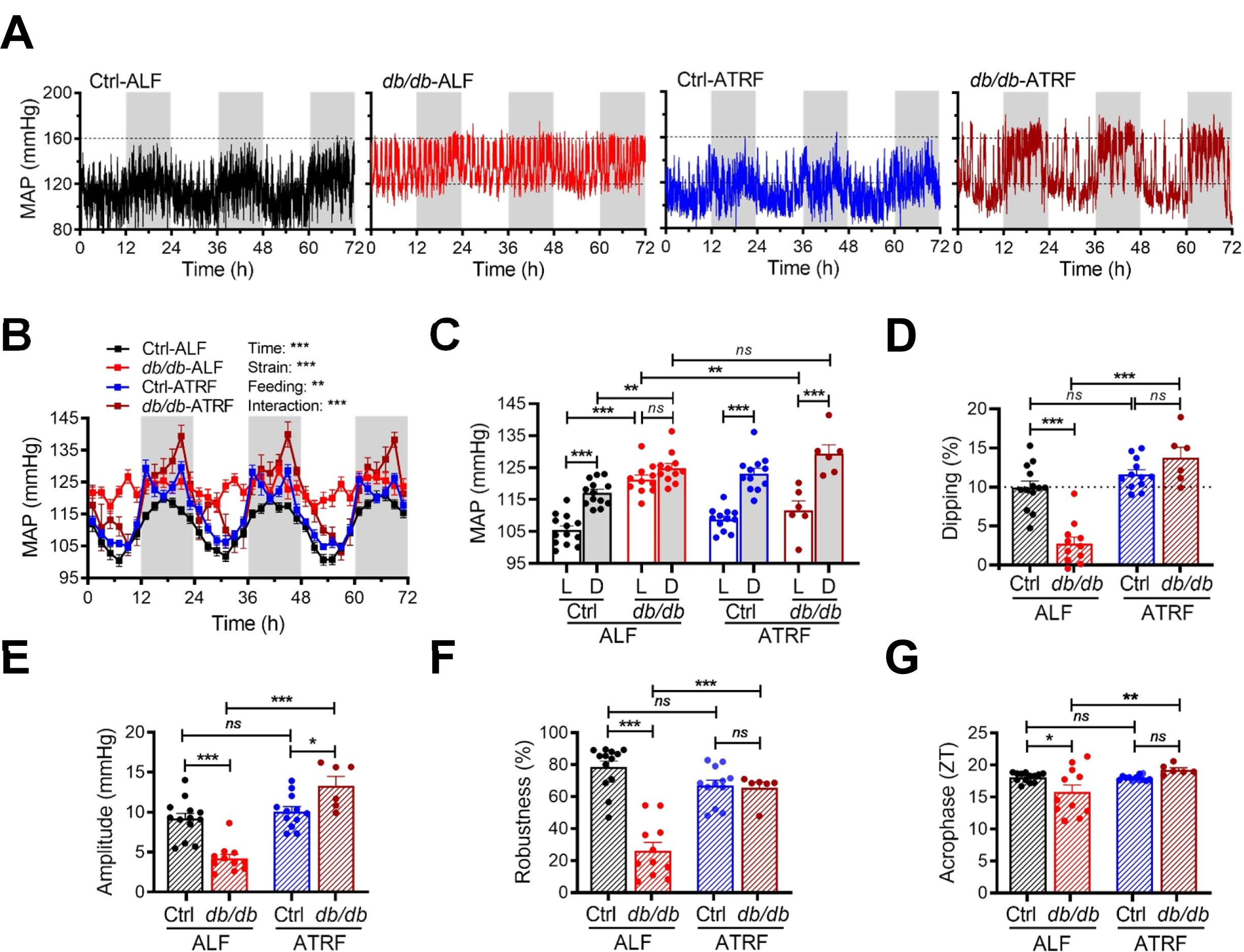
Imposing a food intake circadian rhythm by ATRF prevents *db/db* mice from the development of nondipping blood pressure. Six-week-old *db/db* and control (*db/+*) mice were fed *ad libitum* feeding (ALF) or 8-hour (h) active time-restriction feeding (ATRF) for 10 weeks. (**A**) Representative continuous mean arterial pressure (MAP) recordings by telemetry in 10-second intervals over 72 hours (h) during the light and dark phases, shown in white and gray, respectively. (**B**) Daily profiles of the MAP in 2-h intervals over 72 h during the light and dark phases in Ctrl-ALF (n=13), *db/db*-ALF (n=11), Ctrl-ATRF (n=12), and *db/db*-ATRF (n=6) mice. (**C**) The 12-h average MAP during the light (L) and dark (D) phase. (**D**) MAP dipping (% of MAP decrease during the light phase compared to the dark phase). The dashed line indicates 10% dipping. (**E**−**G**) Amplitude (E), robustness (F), and acrophase (G) of the MAP circadian rhythm. ZT, zeitgeber time (ZT0 = lights-on and ZT12 = lights-off). The data were analyzed by 3-way ANOVA (B and C) and 2-way ANOVA (D−G) with multiple comparisons test. *, p<0.05; **, p<0.01; ***, p<0.001; ns, not significant.

The effect of the imposed diurnal rhythm in food intake on the BP circadian rhythm was further analyzed by cosinor analysis (18). We calculated the amplitude (the half value between the peak and trough of a rhythm), robustness (the strength and stability of a rhythm), and acrophase (the time at which the peak of rhythm occurs) of the MAP circadian rhythm in the same groups of the *db/db* and control mice under ATRF or ALF aforementioned. During ALF, the amplitude, robustness, and acrophase of MAP circadian rhythm were all significantly lower in the *db/db* mice compared to that in the control mice (Figure 2E–2G). In contrast, during ATRF, the amplitude, robustness, and acrophase of MAP circadian rhythm markedly preserved in the *db/db* mice at the levels comparable to control mice (Figure 2E–2G). Importantly, similar effects of ATRF on systolic BP (SBP) and diastolic BP (DBP) circadian rhythms were also found in the same groups of *db/db* and control mice after 10 weeks of ATRF or ALF (Supplemental Figure 3A−3L).

### ATRF restores BP dipping in the *db/db* mice that already have nondipping BP

To investigate whether ATRF can restore BP dipping in *db/db* mice, which is highly relevant to diabetic patients, we investigated the effect of ATRF on BP circadian rhythm in 15-week-old *db/db* mice that already exhibit nondipping BP (15). BP was recorded by telemetry at baseline with ALF and again after 9 days of ATRF. As expected, the *db/db* mice under ALF lost the MAP circadian rhythm (Figure 3A), diurnal MAP variation (Figure 3B), and MAP dipping (Figure 3C). Surprisingly, only 9 days of switching ALF to ATRF sufficiently restored the MAP circadian rhythm (Figure 3A), diurnal MAP variation (Figure 3B), and MAP dipping (Figure 3C). Similar to the effect of ATRF in the prevention studies (Figure 2A–2C), ATRF also selectively dropped the MAP during the light phase but not the dark phase (Figure 3A and Figure 3B), indicating that ATRF restores the BP circadian rhythm through fasting during the light phase in the *db/db* mice. Interestingly, the amplitude and robustness but not the acrophase of MAP circadian rhythm were significantly restored in *db/db*-ATRF mice relative to that in *db/db*-ALF mice (Figure 3D–3F). Moreover, similar effects on the SBP and DBP circadian rhythms were also found in the same groups of *db/db* mice after 9 days of ATRF (Supplemental Figure 4A−4L).

**Figure 3.**
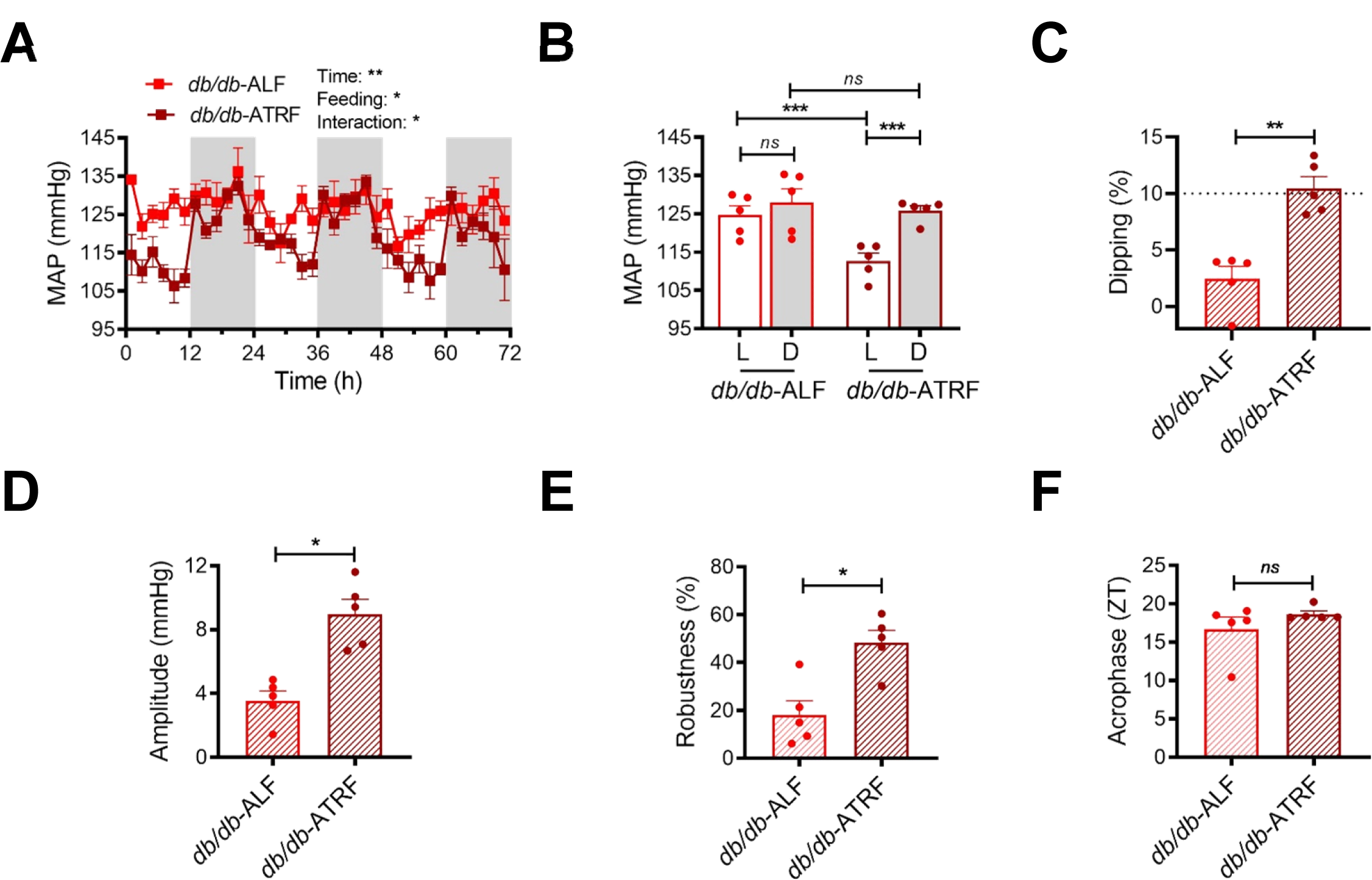
ATRF restores blood pressure dipping in the nondipping *db/db* mice. Blood pressure data were first recorded by telemetry in the 15-week-old *db/db* mice under ALF and then recorded 9 days after ATRF. (**A**) Daily profiles of the mean arterial pressure (MAP) in 2-h intervals over 72 hour (h) during the light and dark phases, shown in white and gray, respectively, in the *db/db*-ALF (n=5) and *db/db*-ATRF (n=5) mice. (**B**) The 12-h average MAP during the light (L) and dark (D) phase. (**C**) MAP dipping. The dashed line indicates 10% dipping. (**D**−**F)** Amplitude (D), robustness (E), and acrophase (F) of MAP circadian rhythm. ZT, zeitgeber time (ZT0 = lights-on and ZT12 = lights-off). The data were analyzed by two-way ANOVA (A and B) with multiple comparisons test and paired t-test (C−F). *, p<0.05; **, p<0.01; ***, p<0.001; ns, not significant.

### ATRF protects *db/db* mice from nondipping BP independent of calorie restriction

Since the mice under ATRF were subjected to 16-hour (h) fasting, they may consume less food than the mice under ALF. As such, they may experience a caloric restriction, which may be responsible for the protective effects of ATRF on the BP circadian rhythm in *db/db* mice. To test this possibility, 6-week-old *db/db* and control mice were fed 8-h ATRF or ALF, and the effects of ATRF on daily food intake, body weight, body composition, blood glucose, and diurnal rhythm in metabolism were determined after 7 to 12 weeks of ATRF or ALF, respectively. Interestingly, ATRF did not affect the daily food intake in control mice but significantly reduced daily food intake in *db/db* mice (Supplemental Figure 5A). Although *db/db*-ATRF mice consumed less food than *db/db*-ALF mice, they had similar body weights and body compositions (Supplemental Figure 5B and 5C). To more accurately determine the effect of ATRF on blood glucose, we measured blood glucose at zeitgeber time (ZT)13 (corresponding to the time after 16-h fasting) and ZT21 (corresponding to the time after 8-h feeding) in *db/db* and control mice under ATRF or ALF. Whereas ATRF did not affect blood glucose at either ZT13 or ZT21 in the control mice, ATRF selectively decreased blood glucose at ZT13 but not ZT21 in the *db/db* mice (Supplemental Figure 5D and 5E). We determined the effect of ATRF on the diurnal rhythm in energy expenditure (EE) and respiratory exchange ratio (RER) in the *db/db* and control mice under ATRF or ALF. ATRF did not affect the EE diurnal rhythm, but markedly enhanced the RER diurnal rhythm in the control mice (Supplemental Figure 6A and 6C). In contrast, ATRF protected the *db/db* mice from the loss of diurnal rhythm in both EE and RER, although the protection was more pronounced for EE than RER (Supplemental Figure 6B and 6D).

To dissect the contribution of calorie restriction *vs.* time restriction to the ATRF protection of BP circadian rhythm, we increased the feeding time from 8 hrs to 12 hrs during the dark-phase. We simultaneously monitored the food intake and BP by BioDAQ and telemetry in the same 14-week-old *db/db* and control mice after 4 days of 12-h ATRF or ALF. Representative food intake recordings illustrated that 12-h ATRF imposed a robust food intake diurnal rhythm on the *db/db* mice with little effect on the control mice (Figure 4A). Quantitative analysis of the food intake recording data showed that 12-h ATRF, however, did not affect the total daily food intake, the number of feeding bouts, duration of feeding bouts, and percent of time spent in feeding bouts in the *db/db* and control mice (Figure 4B and Supplemental Figure 7A−7C). Importantly, similar to 8-h ATRF (Figure 3A and 3B), 12-h ATRFselectively decreased the MAP during the light phase but not during the dark phase, and thus effectively restored the MAP circadian rhythm in the *db/db* mice, including diurnal MAP variation, MAP dipping, amplitude, and robustness (Figure 4C–4H). Since the 12-h ATRF protection of BP circadian rhythm was achieved under iso-caloric feeding, we conclude that ATRF protects *db/db* mice from nondipping BP independent of calorie restriction.

**Figure 4.**
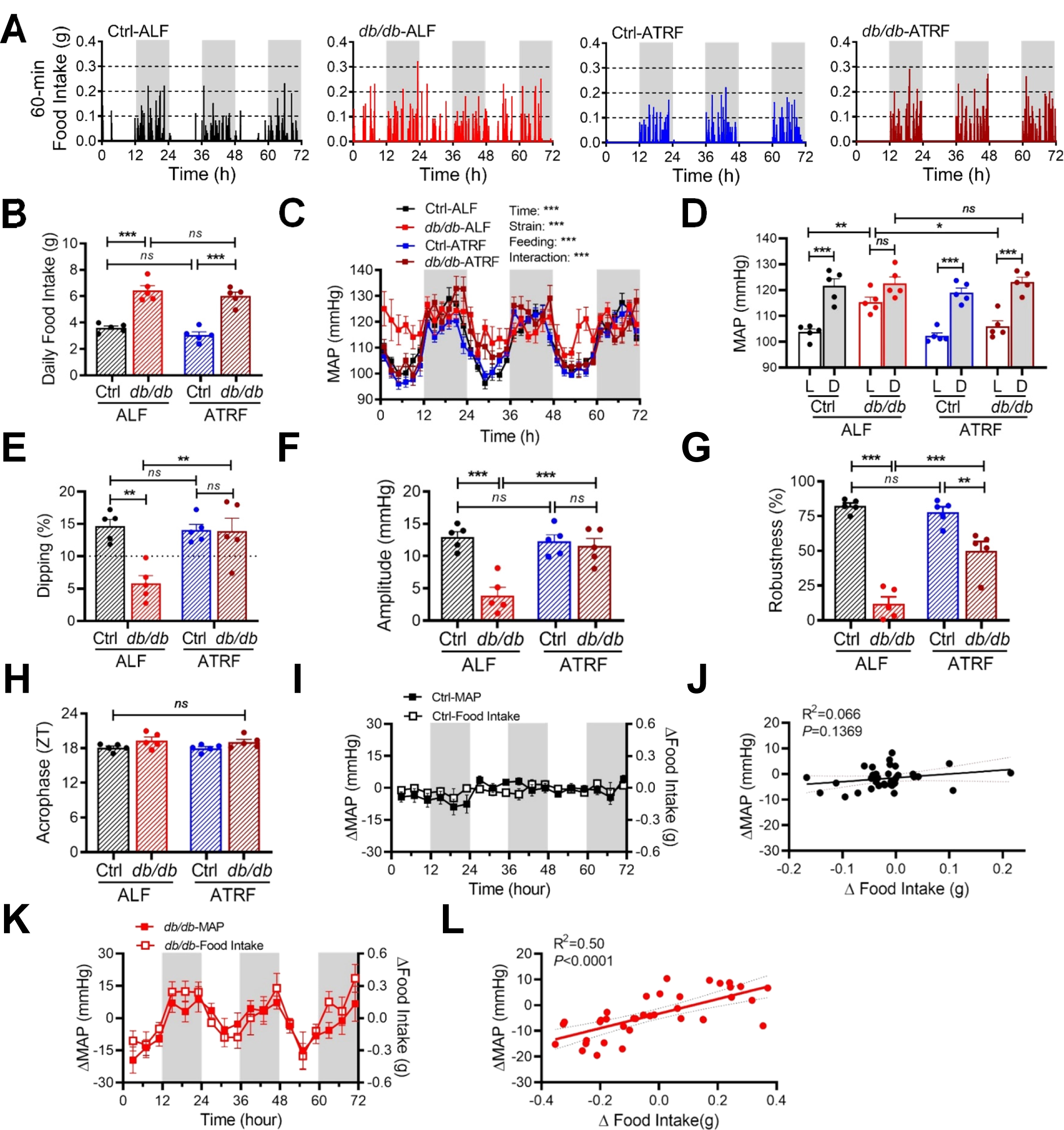
ATRF restores blood pressure dipping in *db/db* mice in the absence of calorie restriction. Food intake and blood pressure were simultaneously recorded by BioDAQ and telemetry in the same 14-week-old *db/db* and control (*db/+*) mice after 4 days of ALF or 12-h ATRF. Five mice in each group. (**A**) Representative Bio-DAQ recordings of daily profiles of the food intake in 1-minute (min) intervals over 72 hours (h) during the light and dark phases, shown in white and gray, respectively. (**B**) Total daily food intake. (**C**) Daily profiles of the mean arterial pressure (MAP) in 2-h intervals during the light and dark phase. (**D**) The 12-h average MAP during the light (L) and dark (D) phase. (**E**) MAP dipping. The dashed line indicates 10% dipping. (**F**−**H**) Amplitude (F), robustness (G), and acrophase (H) of the MAP circadian rhythm. (**I** and **K**) Daily profiles of the changes in MAP (ΔMAP, left y-axis) and food intake (Δfood intake, right y-axis) in 4-h intervals in the control mice (I) and *db/db* mice (K). (**J** and **L**) There is a correlation between ΔMAP and Δfood intake in *db/db* mice (L), but not in control mice (J). The data were analyzed by 2-way ANOVA (B and E−H) and 3-way ANOVA (C and D) with multiple comparisons test and simple linear regression (J and L). *, p<0.05; **, p<0.01; ***, p<0.001; ns, not significant.

Compared with mice under ALF, the *db/db* but not control mice under 12-h ATRF exhibited a selective reduction in food intake and MAP during the light phase than the dark phase (Figure 4A, 4C, and 4D). To investigate whether the reduced consumption of food temporally correlates with the deceased MAP during the light phase, we took advantage of our food intake and BP data that were simultaneously collected from the same *db/db* and control mice. We calculated the change in food intake (Δfood intake) and the change in MAP (ΔMAP) by subtracting the food intake or MAP at corresponding times over 72 hours in *db/db*-ALF and Ctrl-ALF mice from *db/db*-ATRF and Ctrl-ATRF mice, respectively. Strikingly, in the control mice, ATRF induced minimal changes in food intake as well as MAP (Figure 4I), and there was no correlation between Δfood intake and ΔMAP (Figure 4J). In contrast, in the *db/db* mice, ATRF simultaneously induced substantial changes in food intake as well as MAP (Figure 4K), and there was a significant correlation between Δfood intake and ΔMAP (Figure 4L).

### The sympathetic nervous system (SNS) mediates the ATRF protection of BP circadian rhythm in *db/db* mice

To explore the mechanism by which ATRF protects *db/db* mice from nondipping BP, we investigated the potential roles of the SNS as it responds to food intake, exhibits a daily oscillation, and plays a predominant role in the BP circadian rhythm (19, 20). Multiple approaches were taken to investigate this potentially important mechanism.

First, we investigated whether ATRF affects the heart rate (HR) circadian rhythm in *db/db* mice, as HR is regulated by the balance of sympathetic and parasympathetic tone (21). HR circadian rhythm was measured by telemetry in 16-week-old *db/db* and control mice that had been fed ATRF or ALF for 10 weeks. HR circadian rhythm was moderately altered in the *db/db*-ALF mice relative to the Ctrl-ALF mice (Supplemental Figure 8A and 8B). ATRF dramatically decreased the HR during the light phase without affecting the HR during the dark phase and thereby effectively protected the HR circadian rhythm, diurnal HR variation, HR dipping, amplitude, robustness, and acrophase in the *db/db* mice (Supplemental Figure 8A−8F). Similar benefits of ATRF on HR circadian rhythm were also observed in 15-week-old *db/db* mice, whose HR circadian rhythm had already been disrupted (Supplemental Figure 8G−8L).

Second, we investigated whether ATRF affects diurnal variation in HR variability (HRV) in *db/db* mice, as the HRV is considered as an index of the sympathetic and parasympathetic activity (22, 23). The HRV is defined as a physiological variation in the time intervals between heartbeats and can be analyzed by the frequency domain and time domain techniques, including the low-frequency spectral power (LFSP), high-frequency spectral power (HFSP), and root mean square of successive RR interval differences (rMSSD) (22, 23). The effect of ATRF on diurnal variations of HRV was determined in 16-week-old *db/db* and control mice that had been fed ATRF or ALF for 10 weeks. LFSP, HFSP, and rMSSD were all significantly higher during the light phase than during the dark phase in Ctrl-ALF mice (Figure 5A–5C; Supplemental Figures 9A, 9C, and 9E). In contrast, these diurnal variations were abolished in *db/db*-ALF mice (Figure 5A–5C; Supplemental Figures 9B, 9D, and 9F). Interestingly, ATRF significantly increased LFSP, HFSP, and rMSSD during the light phase with little effect during the dark phase, thus preventing *db/db* mice from the loss of the diurnal variations in LFSP, HFSP, and rMSSD (Figure 5A–5C; Supplemental Figure 9A−9F). Similar beneficial effects of ATRF on diurnal variation in HRV were also observed in 15-week-old *db/db* mice that already lost their HRV diurnal variations (Supplemental Figure 10A−10F).

**Figure 5.**
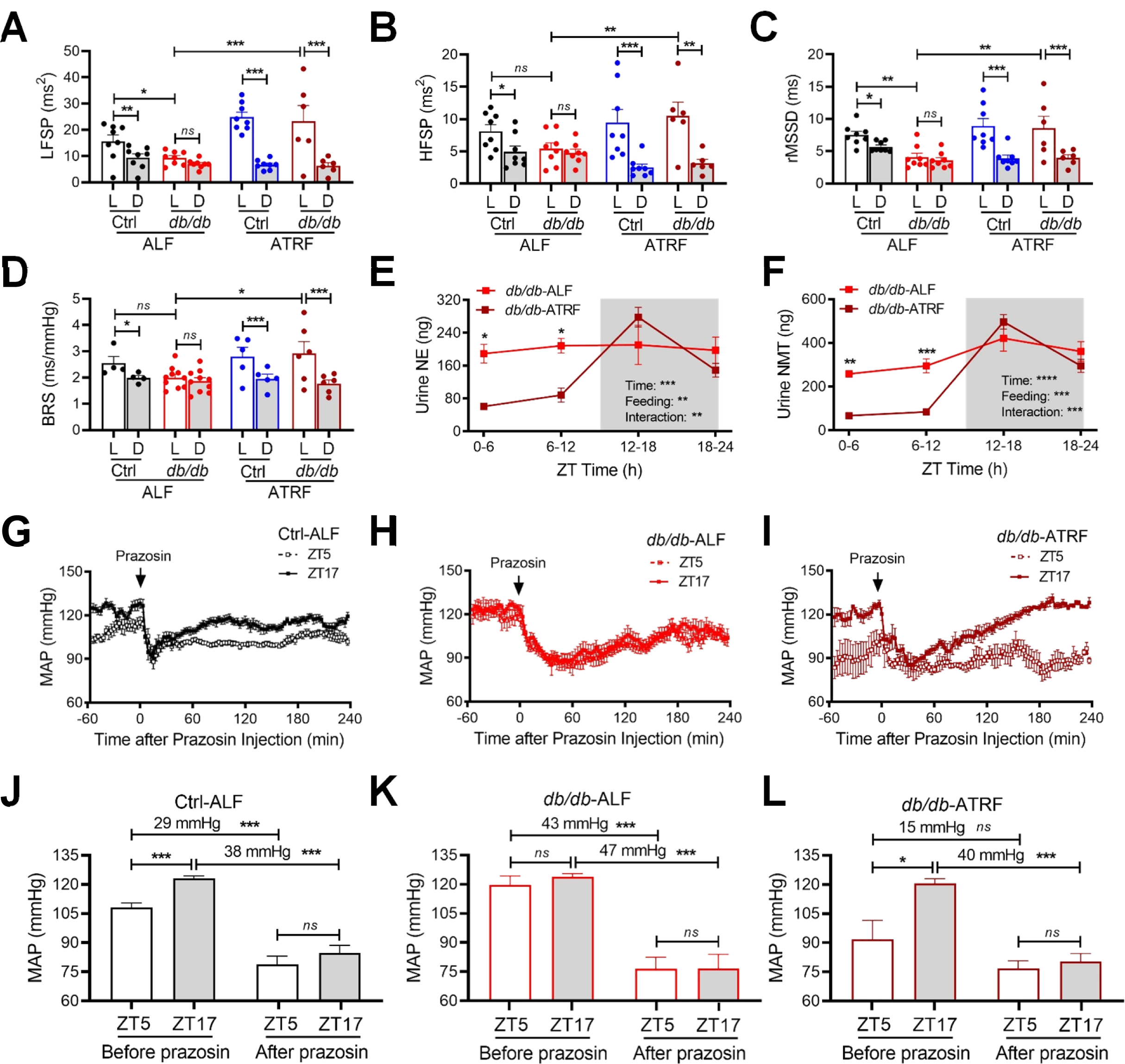
The sympathetic nervous system mediates the ATRF protection of blood pressure circadian rhythm in *db/db* mice. (**A**−**D**) The 12-h average low-frequency spectral power (LFSP; A), high-frequency spectral power (HFSP; B), average root mean square of successive RR interval differences (rMSSD; C) and spontaneous baroreflex sensitivity (BRS; D) during the light (L) and dark (D) phase in 16-week-old *db/db* and control (*db/+*) mice that had been fed ALF or ATRF for 10 weeks. (**E** and **F**) Levels of norepinephrine (NE; E) and its metabolite normetanephrine (NMT; F) in 6-h urine samples collected at ZT0-6, ZT6-12, ZT12-18, and ZT18-24 from 15-week-old *db/db* mice that had been fed ALF or ATRF for 8 weeks (n=4-5). ZT, zeitgeber time (ZT0 = lights-on and ZT12 = lights-off). Levels of NE and NMT are expressed as total contents, which were calculated by concentration x urine volumes. The grey box indicates the dark phase.(**G**−**I**) The 3-minute average MAP response to prazosin, an α1 adrenergic receptor antagonist (1 mg/kg; i.p.), at ZT5 and ZT17 in 18 to 21-week-old Ctrl-ALF (G, n=12), *db/db*-ALF (H, n=9), and *db/db*-ATRF (I, n=4) mice after 14 days of ATRF or ALF. (**J**−**L**) The 1-h average MAP before prazosin injection (baseline) and the lowest MAP after prazosin injection at ZT5 and ZT17 in the Ctrl-ALF (J; n=12), *db/db*-ALF (K; n=9), and *db/db*-ATRF (L; n=4) mice. The data were analyzed by 3-way ANOVA (A−D) and 2-way ANOVA (E−L) with multiple comparisons test. *, p<0.05; **, p<0.01; ***, p<0.001; ns, not significant.

Third, we investigated whether ATRF affects diurnal variation in spontaneous baroreflex sensitivity (BRS) in *db/db* mice, as BRS has been widely used as a cardiac autonomic index (24). The effect of ATRF on BRS was determined by sequence techniques (25) in 16-week-old *db/db* and control mice that had been fed ATRF or ALF for 10 weeks. In accordance with our previous report (16, 26), BRS was higher during the light phase than during the dark-phase in Ctrl-ALF mice (Figure 5D; Supplemental Figure 11A). In contrast, this BRS diurnal variation was lost in *db/db*-ALF mice (Figure 5D; Supplemental Figure 11B). ATRF significantly increased the BRS during the light phase, while having little effect on the BRS during the dark phase, thus protecting *db/db* mice from the loss of the BRS diurnal variation (Figure 5D; Supplemental Figure 11B). ATRF had similar beneficial effects in 15 week-old *db/db* mice that already lost their diurnal variations in BRS (Supplemental Figure 11C and 11D).

Fourth, to more directly investigate whether the SNS is involved in the ATRF protection of BP circadian rhythm in diabetes, we measured diurnal variations in urine norepinephrine (NE) and its metabolite normetanephrine (NMT) in 15-week-old *db/db* mice that had been fed ATRF or ALF for 8 weeks. We were only able to measure NE and NMT in the *db/db* mice due to difficulties in collecting sufficient volumes of urine from the control mice. Nevertheless, our results demonstrated that ATRF potently protected the *db/db* mice from the loss of diurnal variation in both NE and NMT in 6-h urine samples (Figure 5E and 5F; *db/db*-ATRF *vs. db/db*- ALF). Similar with its effects on BP, HR, HRV, and BRS, ATRF also selectively decreased high urine NE and NMT levels during the light phase (ZT0-6 and ZT6-12) but not the dark phase (ZT12-18 and ZT18-24) in the *db/db* mice (Figure 5E and 5F). These results suggest that ATRF may selectively inhibit sympathetic activity during the light phase, thus protecting *db/db* mice from nondipping BP. To verify that the ATRF-induced NE decrease during the light phase in urine samples reflects circulating levels of NE in the *db/db* mice, we measured the plasma NE levels at ZT6 during the light phase in the *db/db* mice under ALF or ATRF. The results confirmed that plasma NE levels were indeed lower during the light phase in *db/db*-ATRF mice than *db/db*-ALF mice (Supplemental Figure 12).

Finally, to explore a potential cause and effect relationship between the sympathetic vascular tone and BP diurnal variation, we determined instant MAP responses to prazosin at ZT5 during the light phase and ZT17 during the dark phase in 18 to 21-week-old *db/db* and control mice after 14 days of ATRF or ALF. Prazosin is a potent α1 adrenergic receptor antagonist that explicitly blocks the effect of the SNS on smooth muscle contraction, thus lowing BP. The average MAP in 1 h before prazosin injection (i.p., 1 mg/kg body weight) and the lowest MAP in 2 hours after prazosin injection were recorded by telemetry, respectively, in Ctrl-ALF, *db/db*- ALF, and *db/db*-ATRF mice. The decrease in MAP between before and after prazosin injection reflected the sympathetic vascular tone at ZT5 or ZT17. In Ctrl-ALF mice, the MAP was higher at ZT17 than ZT5 before prazosin injection, and prazosin reduced MAP to the same value at both times (Figure 5G and 5J). As a result, there was a larger MAP decreasing effect of prazosin at ZT17 than ZT5 (Figure 5G and 5J). This data indicates a higher sympathetic vascular tone during the dark phase than the light phase in the control mice under ALF. In contrast, in *db/db*-ALF mice, the MAP at ZT5 was increased, and the MAP at ZT5 and ZT17 was similar before and after prazosin injection so that prazosin had a similar MAP decreasing effect at both times (Figure 5H and 5K). This data indicates that the sympathetic vascular tone was increased during the light phase in *db/db* mice under ALF, which leads to the loss of the diurnal variation in sympathetic vascular tone. In *db/db*-ATRF mice, the MAP at ZT5 was diminished, and as a result, the diurnal variation in MAP was restored (Figure 5I and 5L). Since the MAP at ZT5 and ZT17 was similar after prazosin injection, there was a larger MAP decreasing effect of prazosin at ZT17 than ZT5 (Figure 5I and 5L). This data indicates that ATRF suppresses the sympathetic vascular tone during the light phase and thereby restored the diurnal variation in MAP in *db/db* mice under ATRF.

In addition to the SNS, we also explored other mechanisms that may play a role in the ATRF protection of BP circadian rhythm in *db/db* mice, including vascular reactivity, adrenal gland hormones, renin-angiotensin-aldosterone system (RAAS), and locomotor activity. Vascular reactivity is defined as the responsiveness of a blood vessel to a specific stimulus. We previously demonstrated that vascular reactivity exhibited a time-of-day variation, which was altered in *db/db* mice and contributed to BP circadian rhythm (27). To explore whether vascular reactivity is involved in the ATRF protection of BP circadian rhythm, we first determined α1a and α1d adrenergic receptor (Adra1a and Adra1d) mRNA daily oscillations in the mesenteric arteries of 21-week-old *db/db* and control mice that had been fed ALF or ATRF for 4 weeks.

As shown in Supplement Figure 13A and 13B, ATRF did not affect Adra1a and Adra1d mRNA daily oscillations in *db/db* or control mice. To more directly explore the role of vascular reactivity in the ATRF protection of BP circadian rhythm, we determined the instant pressor responses to an intravenous bolus injection of phenylephrine (PE) or angiotensin II (Ang II) at ZT5 and ZT17 in *db/db* mice after 9 days of ATRF. If ATRF protects BP circadian rhythm via vascular reactivity, we expected that ATRF would selectively inhibit vascular reactivity at ZT5 during the light phase. In contrast to our expectation, there was no significant difference in the pressor responses to PE or Ang II between ZT5 and ZT17 in *db/db*-ATRF mice (Supplement Figure 13C and 13D), indicating that vascular reactivity is unlikely involved in the ATRF protection of BP circadian rhythm.

Adrenal gland hormones, including epinephrine (EPI), aldosterone (ALDO), and corticosterone (CORT), are regulated by the SNS and are well recognized for their pivotal roles in BP and glucose homeostasis (28). To explore whether these adrenal gland hormones are involved in the ATRF protection of BP circadian rhythm, we determined EPI, ALDO, and CORT diurnal variations in the same 6-h urine samples that we used for the measurement of NE and NMT diurnal variation in *db/db* and control mice under ATRF or ALF. In contrast to NE and NMT (Figure 5E and 5F), EPI, ALDO, and CORT possessed clear-cut diurnal variation in *db/db*-ALF mice (Supplemental Figure 14A−14C). Similar to its effect on NE and NMT (Figure 5E and 5F), ATRF suppressed ALDO and CORT during the light phase (ZT6-12), thus improving their diurnal variations (Supplemental Figure 14B−14C). On the contrary, ATRF raised EPI during the dark phase (ZT12-18), thus enhancing its diurnal variation (Supplemental Figure 14A).

The RAAS is primarily comprised of angiotensinogen (Agt), renin (Ren1), angiotensin-converting enzyme (Ace), Ace2, and angiotensin II receptor 1a (Agtr1a). Accumulating evidence suggests that activation of the RAAS under various pathological conditions, including type 2 diabetes, contributes to the disruption of BP circadian rhythm via the SNS (17, 29). To explore whether the RAAS is involved in the ATRF protection of BP circadian rhythm in *db/db* mice, we determined the daily oscillation of Agt, Ren1, Ace, Ace2, and Agtr1a mRNAs in the liver or kidney from 21-week-old *db/db* and control mice that had been fed ATRF or ALF for 4 weeks. There were some differences in Agt, Ren1, Ace, Ace2, and Agtr1a mRNA daily oscillations between *db/db* and control mice, but no significant effect of ATRF was found in *db/db* mice (Supplement Figure 15A−15F).

Locomotor activity and BP are synchronized by the clock genes in the SCN and exhibit similar circadian rhythms under physiological conditions (26). We have previously reported that the circadian rhythms in locomotor activity and BP are severely suppressed in *db/db* mice (15). To determine whether ATRF protects BP circadian rhythm through locomotor activity, we measured locomotor activity by telemetry in the same *db/db* and control mice under ATRF or ALF in which we measured BP (Figure 2). Consistent with our previous report (15), the *db/db* mice under ALF showed no locomotor activity circadian rhythm (Supplemental Figures 16A and 16B). In contrast to its potent protection on the BP circadian rhythm in the *db/db* mice (Figures 2), ATRF but did not lead to a robust locomotor activity circadian rhythm in the *db/db* mice (Supplemental Figure 16A−16F). In 15-weeks-old *db/db* mice that already had lost their locomotor activity circadian rhythm, ATRF induced a brief increase in locomotor activity at the onset of the dark phase but failed to restore the locomotor activity circadian rhythm (Supplemental Figure 16G−16L).

### Identification of BMAL1 as a key molecular player in the ATRF protection of BP circadian rhythm in *db/db* mice

Circadian rhythms are controlled at the molecular level by interacting transcriptional-translational feedback loops composed of a set of clock genes, which are expressed in cells throughout the body (30). To investigate whether the clock genes participate in the ATRF protection of BP circadian rhythm, we determined the daily transcript oscillations of a panel of clock genes, including Bmal1 (also known as Arntl3 in mouse and MOP3 in humans), Per 1 (period 1), Per2 (period 2), Cry 1(cryptochrome 1), Cry2 (cryptochrome 2), Clock, Rev-erbα (also known as nuclear receptor subfamily 1, group D, member 1 [Nr1d1)], and Rorc (RAR-related orphan receptor gamma), in the liver, kidney, heart, adrenal gland, and mesenteric arteries from *db/db* and control mice. We chose the liver because clock genes in the liver are extensively studied and are well documented for their robust and prompt responses to feeding (31). We chose the kidney, heart, adrenal gland, and mesentery arteries because these tissues are critical for BP homeostasis and circadian rhythm (32). We did not choose the suprachiasmatic nucleus (SCN), the master circadian pacemaker that responds to environmental light, because previous studies have shown that that the SCN does not respond to time-restricted feeding (31).

The daily mRNA oscillation of Bmal1, Per1, Per2, Cry, Cry2, Clock, Rev-erbα, and Rorc were determined by real-time PCR in the liver, kidney, heart, adrenal gland, and mesentery arteries collected at ZT5, ZT11, ZT17, and ZT23 from 21-week-old *db/db* and control mice that had been fed ATRF or ALF for 4 weeks. Consistent with our previous report (27), the daily mRNA oscillation of Bmal1, Per1, Per2, Cry, Cry2, Clock, Rev-erbα, and Rorc were all significantly altered in the *db/db* mice compared to the control mice under ALF (Figure 6A–6E and Supplemental Figure 17−21). Importantly, ATRF was capable of restoring the daily mRNA oscillation of Bmal1, Per1, Per2, Cry, Cry2, Clock, Rev-erbα, and Rorc in the *db/db* mice in a gene and tissue-specific manner (Figure 6A–6E and Supplemental Figure 17−21). Of particular interest is that Bmal1, among all genes examined, was most dramatically and consistently restored by ATRF in all tissues examined (Figure 6A–6E *vs.* Supplemental Figure 17 and 21). In Ctrl-ALF mice, Bmal1 mRNA oscillated with the peak at ZT23 and trough at ZT11 in all tissues examined (Figure 6A–6E). In contrast, in *db/db*-ALF mice, Bmal1 mRNA oscillated with the peak at ZT17 and trough at ZT5 in all tissues examined except for the mesentery arteries in which Bmal1 daily oscillation was suppressed (Figure 6A–6E). Regardless of this difference, The timing of the Bmal1 mRNA daily oscillation was notably restored in all tissues examined in *db/db*-ATRF mice compared to *db/db*-ALF and/or Ctrl-ALF mice (Figure 6A–6E).

**Figure 6.**
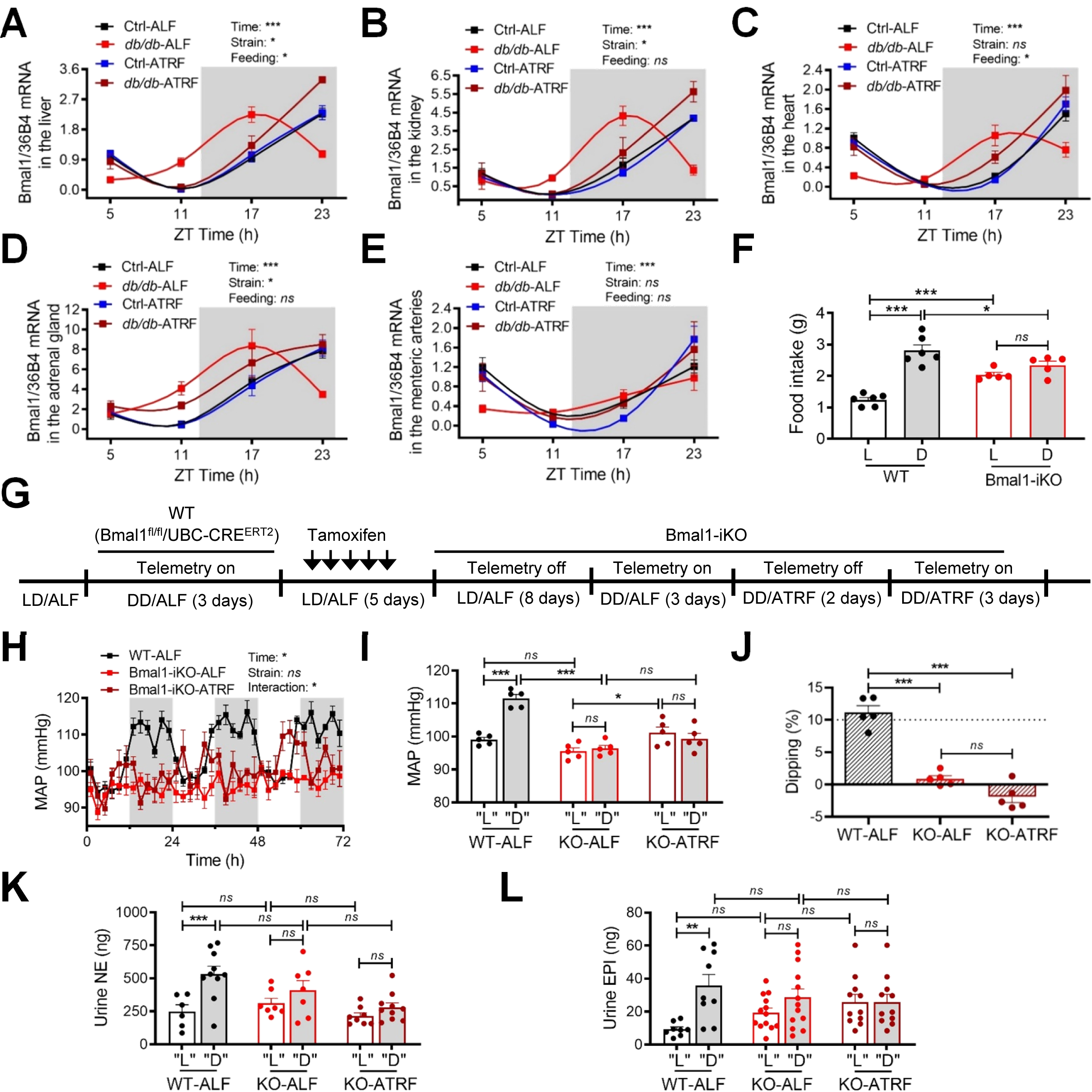
Identification of BMAL1 as a key molecular player in the ATRF protection of BP circadian rhythm in *db/db* mice. (**A**−**E**) Bmal1 mRNA daily oscillation at zeitgeber time (ZT)5, ZT11, ZT17, and ZT23 in the liver (A), kidney (B), heart (C), adrenal gland (D), and mesenteric arteries (E) from 21-week-old Ctrl-ALF (n=6-7), Ctrl-ATRF (n=4-5), *db/db*-ALF (n=4-5), and *db/db*-ATRF (n=3-5) mice that had been fed ALF or ATRF for 4 weeks. Bmal1 mRNA levels were expressed as the ratio of Bmal1 to 36B4 relative to that in the Ctrl-ALF mice at ZT5. The grey box indicates the dark phase. (**F**) The 12-h food intake during the light (L) and dark (D) phase in 16-week-old Bmal1-iKO and WT control mice under ALF. (**G**) Scheme of the experimental design. All experiments were performed under constant darkness except the days for tamoxifen injection and recovery. LD, light-dark cycle. DD, constant darkness. (**H**) Daily profiles of mean arterial pressure (MAP) in 2-h intervals over 72 h in WT-ALF, Bmal1-iKO-ALF, and Bmal1-iKO-ATRF mice. Five mice in each group. (**I**) The 12-h average MAP during the subjective light (“L”) and subjective dark (“D”) phases. (**J**) MAP dipping. The dashed line indicates 10% dipping. (**K** and **L**) Total contents of 12-h urine norepinephrine (NE; K) and epinephrine (EPI; L) during the subjective light (“L”) and subjective dark (“D”) phases. The data were analyzed by 2-way ANOVA (A−I, K and L) and 1-way ANOVA (J) with multiple comparisons test. **, p<0.01; **, p<0.01; ***, p<0.001; ns, not significant.

Among the canonical clock genes, BMAL1 is the only one whose deletion results in immediate and complete loss of circadian rhythmicity (33), including loss of the BP circadian rhythm (34, 35). To investigate whether BMAL1 is involved in the ATRF protection of BP circadian rhythm, we generated a tamoxifen-inducible global Bmal1 knockout mouse model (Bmal1-iKO) by crossing the Bmal1 flox mice (36) with the UBC-Cre^ERT2^ mice that ubiquitously express Cre enzyme (37). Consistent with the previous report (35), Bmal1-iKO mice were viable, fertile, and grossly normal. By genomic PCR, we demonstrated that the Cre-mediated chromosome recombination was only detectable in Bmal1-iKO mice (Bmal1^fl/fl^/UBC-Cre^ERT2^) but not in WT mice (Bmal1^fl/fl^; Supplemental Figure 22A). By Western blot, we illustrated that BMAL1 protein was barely detectable in Bmal1-iKO mice compared to its abundant expression in WT mice in the liver, heart, kidney, and adrenal gland (Supplemental Figure 22B).

To investigate whether the global ablation of Bmal1 affects the food intake diurnal rhythm, we measured food intake by indirect calorimetry in 16-week-old Bmal1-iKO and WT mice under ALF. In contrast to WT control mice, Bmal1-iKO mice did not exhibit the food intake diurnal rhythm with food consumed almost equally during both phases (Figure 6F and Supplemental Figure 23A and 23B). We then determined whether imposing a food intake diurnal rhythm by ATRF can similarily protect the BP circadian rhythm in Bmal1-iKO mice as it protects *db/db* mice from nondipping BP. To minimize the individual animal variability, we recorded BP in the same single-housed mouse before and after tamoxifen injection (Figure 6G). Since global Bmal1 knockout mice have defective inner retinal electrical responses to light (36), we recorded BP under the constant darkness (DD) in 16-week-old Bmal1-iKO and WT mice under ATRF or ALF to eliminate potential confounding effects (Figure 6G).

In accordance with our previous report (26), WT-ALF mice (before tamoxifen injection) exhibited a robust MAP circadian rhythm with a significantly higher MAP during the subjective dark phase (the 12-h period corresponding to the previous dark phase in the light (L)/dark (D) cycle) than that during the subjective light phase (the 12-h period corresponding to the previous light phase in the L/D cycle; Figure 6H and 6I). In contrast, Bmal-iKO-ALF mice (after tamoxifen injection) had a substantial MAP drop during the subjective dark phase, which led to the complete loss in the MAP circadian rhythm, MAP diurnal variation, and MAP dipping (Figure 6H–6J). Cosinor analysis revealed of the MAP circadian rhythm showed a dramatic reduction in the amplitude, robustness, and acrophase of MAP circadian rhythm in Bmal1-iKO mice relative to the control mice (Supplemental Figure 24A−24C). Importantly, in contrast to its striking protective effect on BP circadian rhythm in *db/db* mice (Figure 2–4), ATRF failed to restore the MAP circadian rhythm, MAP diurnal variation, and MAP dipping in BMAL1-iKO mice (Figure 6H–6J). ATRF also was unable to restore the amplitude and robustness of the MAP circadian rhythm in BMAL1-iKO mice (Supplemental Figures 24A and 24B), indicating that BMAL1 is essential for the ATRF protection of BP circadian rhythm.

Finally, we investigated the possible role of the SNS in linking ATRF/BMAL1 to the BP circadian rhythm. NE and EPI were measured by ELISA in 12-h urine samples collected from 16-week-old Bmal1-iKO and WT mice that had been fed ATRF or ALF under constant darkness for 6 days. As expected, in WT-ALF mice, both NE and EPI were higher during the subjective dark phase than the subjective light phase (Figure 6K and 6L), which correlated with a higher MAP during the subjective dark phase than the subjective light phase (Figure 6H and 6I). Surprisingly, in contrast to a crystal-clear suppression of NE and EPI in a constitutive BMAL1 knockout mouse model (34), both NE and EPI were not significantly changed in BMAL1-iKO-ALF mice (Figure 6K and 6L). Nevertheless, in BMAL1-iKO-ALF mice, the diurnal rhythms in NE and EPI were abolished (Figure 6K and 6L), which correlated with the loss of the BP circadian rhythm (Figure 6H and 6I). In contrast to its dramatic restoration of NE (Figure 5E) and EPI (Supplemental Figure 14A) and BP circadian rhythms in *db/db* mice, ATRF was unable to restore diurnal rhythms in NE and EPI (Figure 6K and 6L) or BP in BMAL1-iKO-ALF mice (Figure 6H and 6I).

## Discussion

The major findings of the current study include 1) loss of the food intake diurnal rhythm coincides with nondipping BP in diabetic *db/db* mice, 2) imposing a food intake diurnal rhythm with fasting during the light phase (i.e., ATRF) prevents *db/db* mice from the development of nondipping BP, 3) ATRF, independent of calorie restriction, restores BP dipping in *db/db* mice that already have nondipping BP, 4) the SNS mediates the ATRF protection of BP circadian rhythm in *db/db* mice, 5) Identification of BMAL1 as a key molecular player in the ATRF protection of BP circadian rhythm in *db/db* mice.

What is the significance of our findings? There is a growing body of literature demonstrating that the timing of food intake is critical for diet-induced metabolic disorders (9-13). However, only sparse information about the effect of the timing of food intake on BP circadian rhythm is presently available. It has been shown that changing the time when pellet food was presented to rabbits or dogs immediately shifted the peak of the BP circadian rhythm (38, 39). It has also been shown that rats receiving feeding for 1 h only during the light phase rather than the dark phase exhibited a suppressed BP circadian rhythm (40). These studies suggest that the timing of food intake can influence BP circadian rhythm in nondiabetic healthy animals. However, it is unknown whether nondipping BP in diabetes, which is a serious risk factor for adverse cardiovascular consequences, is causally associated with altered daily timing of food intake.

The current study demonstrated that the loss of food intake diurnal rhythm coincides with nondipping BP in type 2 diabetic *db/db* mice (Figure 1, Figure 2A, and Figure 2B). More importantly, the current study demonstrated for the first time that imposing a food intake diurnal rhythm by ATRF not only dramatically prevented *db/db* mice from the development of nondipping BP (Figure 2 and Supplemental Figure 3) but also effectively restored BP dipping in *db/db* mice that already have nondipping BP (Figure 3, Figure 4, and Supplemental Figure 4). The results from the current study suggest that the loss of food intake diurnal rhythm contributes to the etiology and development of nondipping BP in diabetes, and the lifestyle changes (i.e., limiting eating to the right time of day) may effectively improve nondipping BP in diabetes.

The ATRF used in the current study differs from intermittent fasting or periodic fasting that lately has become very popular in humans (41). ATRF, intermittent fasting, and periodic fasting all impose a cycle consisting of fasting periods alternating with nonfasting periods and are all effective in protection against obesity, diabetes, cancers, heart disease, asthma, rheumatoid arthritis, neurodegeneration, and hypertension (9, 41). However, the ATRF used in the current study imposes the fasting and feeding cycle within one 24-h day without an overt attempt to reduce calorie intake, and importantly, the feeding phase aligns with the dark phase when most of the food is normally consumed in nocturnally active rodents. In contrast, the intermittent fasting imposes fasting for 1 day every other day or 2 days a week, and periodic fasting imposes fasting three days or longer every 2 or more weeks, both of which have 20- 40% of calorie restriction (41). Given that BP exhibits a circadian rhythm, it is tempting to speculate that ATRF, but not intermittent fasting or periodic fasting, can effectively protect against nondipping BP in diabetes. The current study supports this speculation. However, further research is needed to address this issue.

The ATRF used in the current study also differs from the bedtime ingestion of anti-hypertension medications, which has been shown to be more effective than traditionally recommended morning ingestion of anti-hypertension medicines (42). Compared with morning ingestion of anti-hypertension medications, bedtime ingestion of anti-hypertension medications not only improves BP control and nondipping BP but also significantly reduces the risk of new- onset type 2 diabetes and cardiovascular events (42, 43). However, it is unlikely that bedtime ingestion of anti-hypertension medications is effective for the treatment of metabolic disorders that commonly co-exist with hypertension in type 2 diabetic patients. It is also unclear whether bedtime ingestion of anti-hypertension medications is effective in the prevention of the development of metabolic disorders and nondipping BP in people with obesity and prediabetes. In contrast, the ATRF used in the current study not only effectively restores the BP circadian rhythm in type 2 diabetic mice that have already developed nondipping BP (Figures 3 and 4) but also dramatically prevents the development of nondipping BP (Figure 2) and metabolic disorder (Supplemental Figure 6) in type 2 diabetic mice (Figure 2). A recent small but rigorously controlled proof-of-concept clinical trial reported that early time-restricted feeding, independent of body weight loss, effectively improved insulin sensitivity and oxidative stress and dramatically lowered SBP and DBP in men with prediabetes (44). Although this clinical trial did not monitor BP circadian rhythm, it suggests the feasibility of ATRF in the prevention of nondipping BP in humans. However, large clinical trials are needed to investigate further the efficacy of this nonpharmacological, chrono-nutritional, and low-cost approach for restoration and maintenance of a healthy BP rhythm in diabetic patients.

How does ATRF protect BP circadian rhythm in diabetes? One of the intriguing findings from the current study is that imposing food intake diurnal rhythm by ATRF selectively lowed BP during the light or fasting phase but not the dark or nonfasting phase in *db/db* mice (Figure 2B and 2C, Figure 3A and 3B, Figure 4F and 4G). While the underlying mechanism remains elusive, previous studies have shown that nondipping BP was associated with increased sympathetic vascular tone in *db/db* mice (15, 17), and fasting was capable of potently suppressing sympathetic activity in nondiabetic rats (20, 45). These studies suggest that ATRF may selectively decrease sympathetic vascular tone during the light phase via fasting in *db/db* mice and thereby protect their BP circadian rhythm. Several lines of evidence from the current study support this potential mechanism. First, ATRF selectively decreased HR during the light phase but not the dark phase in *db/db* mice (Supplemental Figure 8). Second, ATRF selectively increased the HRV (LFSP, HFSP, and rMSSD) during the light phase but not the dark phase in *db/db* mice (Figure 5A–5C and Supplemental Figure 9 and 10). Third, ATRF selectively increased the BRS during the light phase but not the dark phase in *db/db* mice (Figure 5D and Supplemental Figure 11). Since the HR, HRV, and BRS reflect the balance of sympathetic and parasympathetic activity (22, 23), these findings suggest that ATRF protects BP circadian rhythm in *db/db* mice by inhibiting the sympathetic activity or activating parasympathetic vascular tone during the light phase via fasting in *db/db* mice. Fourth, perhaps the most important, ATRF selectively decreased sympathetic neurotransmitter NE and its metabolite NMT during the light phase but not the dark phase in *db/db* mice (Figure 5E and 5F; Supplemental Figure 12). Finally, the selective effect of ATRF on BP during the light phase was sensitive to α1 adrenergic receptor antagonist prazosin in *db/db* mice (Figure 5G–5L). Collectively, these data suggest that ATRF inhibits sympathetic vascular tone during the light phase via fasting, thus protecting the BP circadian rhythm in *db/db* mice.

In addition to the SNS, accumulating evidence suggests that many additional mechanisms may also mediate the ATRF protection of BP circadian rhythm in *db/db* mice. These potential mechanisms include but not limited to vascular reactivity, adrenal gland hormones, RAAS, locomotor activity, sleep-wake cycle, and gut microbiota. While the results from the current study do not support the role of vascular reactivity and locomotor activity in the ATRF protection of BP circadian rhythm in *db/db* mice (Supplemental Figures 13 and 16), the following mechanisms need to be investigated further. First, although we did not find that ATRF affects the daily mRNA oscillation of Agt, Ren1, Ace1, Ace2, and Agtr1a in the liver or kidney in *db/db* mice (Supplemental Figure 15), the role of the RAAS in the ATRF protection of BP circadian rhythm in diabetes cannot be excluded. It has been shown that chronic treatment of *db/db* mice with the Ang II antagonist losartan or the ACE inhibitor enalapril blocked the increase in MAP and improved autonomic regulation (17, 29). It also has been shown that feeding time has a marked influence on the chronobiology of renin cascade, urinary electrolytes, and BP (39). Consistent with these findings, the current study demonstrated that ATRF selectively diminished urine aldosterone during the light phase but not the dark phase in *db/db* mice (Supplemental Figure 14B). Second, we have shown in other studies that ATRF can restore a normal daily sleep-wake cycle in *db/db* mice (46). Since the role of the sleep-wake cycle in BP circadian rhythm is well recognized (47), ATRF may improve the sleep-wake cycle and thereby protect the BP circadian rhythm in *db/db* mice. Third, several recent studies indicated that gut microbiota is involved in BP control and hypertension (48). Moreover, the gut microbiome is highly dynamic and exhibits daily cyclical fluctuations in composition, which can be dramatically affected by ATRF and thus links to host circadian clock and metabolism (49).

How does BMAL1 coordinate with the SNS to participate in the ATRF protection of BP circadian rhythm in diabetes? It has been shown that the mice lacking Bmal1 exhibit severe hypotension and a complete loss in the BP circadian rhythm (34). While multiple mechanisms likely contribute to these phenotypes, it has been shown that the plasma catecholamines (NE and EPI) and the enzymes accounting for their biosynthesis and degradation in the heart and adrenal gland were markedly suppressed in the mice lacking Bmal1 (34). These data suggest that the BMAL1 can regulate the biosynthesis or degradation of catecholamines. Consistent with these findings, it has been shown that BMAL1 can bind to and activate the promoter of monoamine oxidase, an enzyme crucial for NE degradation (50). We and others previously reported that Bmal1 mRNA daily oscillation was altered in various types of tissues from *db/db* mice (15, 27, 51). In line with these reports, the current study further demonstrated that ATRF effectively restored Bmal1 mRNA daily oscillation in the liver, kidney, heart, adrenal gland, and mesenteric arteries from *db/db* mice (Figure 6A–6E). It has been shown that nuclear BMAL1 protein, but not total BMAL1 protein, was suppressed by ATRF in the mouse liver during the fasting light phase (52), which coincides with the current finding that ATRF selectively reduced BP and NE during the fasting light phase (Figure 2-4 and Figure 5E). Given the predominant role of the SNS in the BP circadian rhythm, these data, taken together, suggest a potential mechanism by which ATRF inhibits BMAL1 and then attenuates the sympathetic vascular tone via fasting during the light phase, thus protecting the BP circadian rhythm in diabetic mice. The current findings that ATRF fails to restore diurnal catecholamine variations and BP circadian rhythms in the mice lacking Bmal1 (Figure 6G–6L) further supports this putative mechanism.

Nevertheless, it should be pointed out that in addition to the SNS, BMAL1 may mediate the BP circadian rhythm through other mechanisms, including the RAAS, sleep-wake cycle, and gut microbiota, which will need to be investigated further.

There are some limitations in the current studies that need to be addressed in future studies. First, the molecular mechanisms by which ATRF modulates the SNS via BMAL1 remain to be investigated. Second, whether the ATRF used in the current study is also effective in the protection of BP circadian rhythm in females as well as males, and other diabetic and nondipping BP animal models remains to be elucidated. Third, whether the ATRF used in the current study also effectively protects target organ damage that is associated with nondipping BP in diabetic patients remains to be explored.

In summary, the current study demonstrated for the first time that ATRF, independent of calorie restriction, effectively protects diabetic *db/db* mice from nondipping BP. Mechanistically, we unveiled that imposing fasting during the light phase by ATRF inhibits sympathetic activity, thus protects BP circadian rhythm in *db/db* mice. Moreover, we identified BMAL1 as a potentially key molecular player that links ATRF, the sympathetic vascular tone, and BP circadian rhythm.

Importantly, the current study suggests that people with prediabetes or patients with diabetes can improve their BP circadian rhythm by lifestyle changes (i.e., eating only at the right time), which can potentially improve their health outcomes.

## Methods

### Animals

The following mice were purchased from the Jackson Laboratories (Bar Harbor, ME): *db/db* and nondiabetic *db/+* mice (Stock No.: 000642), Bmal1^*flox/flox*^ mice (Storch et al., 2007), (Stock No.: 007668), and UBC-Cre-ERT2^+/-^ mice (Stock No.: 007001). The inducible global Bmal1 knockout mice (Bmal1-iKO) were generated by crossing the Bmal1^*flox/flox*^ with the UBC-Cre-ERT2^+/-^ mice and IP injection of 100 µl tamoxifen (20 mg/ml in sesame oil, q.d. for 5 consecutive days). All mice were fed with chow diet with free access to water, and kept in a light-tight box under a 12:12 light: dark cycle unless indicated otherwise. Male mice were used in the current study. All animal procedures were approved by the Institutional Animal Care and Use Committee of the University of Kentucky.

### Feeding schedules

The mice were fed ALF (ad libitum), 8-h ATRF (8 hours of food access from ZT13 to ZT21 [ZT0 is the time of lights-on, and ZT12 is the time of lights-off]), or 12-h ATRF (12 hours of food access from ZT12 to ZT24) for the periods as described in the result and figure legends.

### Food intake diurnal rhythm measurement

The food intake diurnal rhythm was comprehensively characterized by using a metabolic chamber (indirect gas calorimetry LabMaster system, TSE System, Bad Homburg, Germany) or a BioDAQ system (Research Diet, New Brunswick, NJ). The mice were acclimated to the systems for at least 7 days; then, the food intake was quantified continuously for a minimal of 3 consecutive days.

### Telemetric measurement of the circadian rhythms of BP, heart rate, and locomotor activity

The mice were chronically instrumented in the left common carotid artery with a telemetry probe (TA11PA-C10, Data Sciences International, St. Paul, MN, USA) as described previously (15, 16, 26). After 7-10 days of recovery, systolic BP (SBP), diastolic BP (DBP), heart rate (HR), and locomotor activity were recorded for at least 3 consecutive days in home-caged conscious free-moving mice. The telemetric data were analyzed by Dataquest A.R.T. software (Data Sciences International [DSI], St. Paul, MN).

### Metabolic characterizations of the animals

Body composition (lean mass and fat mass) was assessed by NMR spectroscopy (Echo MRITM-100H, Houston, TX). Nonfasting blood glucose was measured by a StatStrip® XepressTM glucometer (NOVA® biomedical, Waltham, MA, USA). Respiratory exchange ratio (RER) and energy expenditure (EE) were measured using the metabolic chamber. Oxygen consumption and carbon dioxide production in the metabolic chambers were measured every 30 min for 3 consecutive days. RER and EE were calculated using the accompanying TSE PhenoMaster software.

### Urinary and plasma catecholamines, aldosterone, and corticosterone measurement

The mice were acclimated in the metabolic cage (Tecniplast, S.p.A.) for about 12 hours, and then urine samples were collected during 6-h intervals from ZT0 to 6, ZT6 to 12, ZT12 to 18, and ZT18 to 24. Plasma samples were collected at ZT6 during the light phase in *db/db* mice. The epinephrine, norepinephrine, and normetanephrine in urine or plasma were determined by the ELISA kits (Abnova, Taipei, Taiwan). The urinary aldosterone and corticosterone were determined by the ELISA kits from Enzo Life Sciences, Inc. (Farmingdale, NY) and Arbor Assay (Ann Arbor, MI), respectively. Total contents of epinephrine, norepinephrine, normetanephrine, aldosterone, and corticosterone in the 6-h urine samples were calculated by concentrations × urine volumes.

### Heart rate variability analysis

Heart rate variability was analyzed from the telemetry data by frequency domain and time domain methods using Ponemah Software (Data Sciences International; St. Paul, MN, version 6.32). For frequency domain analysis, 2 min segments at 20 min interval over 72 hours were selected and scanned to ensure they were free of artifacts. Each segment was interpolated to 20 Hz using the quadratic method. Subsequently, the data were subdivided into 50 overlapping series and computed by Fast Fourier Transform using the Hanning window method. The cut-off frequency range for low-frequency (LF) was 0.15-0.6 Hz, optimized by Baudrie’s group (22). The high-frequency (HF) range was 1.5-4 Hz. For time-domain analysis, 5 min segments over 72 hours were calculated, and the root-mean-square successive beat-to-beat difference (rMSSD) was plotted as the marker of parasympathetic heart rate control. For both the frequency and time domain data, the heart rate variability was averaged in each correspondent hour over 3 days to generate the 24-h heart rate variability profile.

### Baroreflex sensitivity analysis

The spontaneous baroreflex sensitivity was analyzed by sequence techniques using the Hemolab software (http://www.haraldstauss.com/HemoLab/HemoLab.html) <u>as previously described</u> (26). For every 1-h data, the software finds the sequences in which the systolic arterial pressure and pulse interval were positively correlated (r^2^>0.80) and counts those that contain at least four consecutive sequences as an effective Baroreflex. Baroreflex sensitivity was calculated as the average slope of the systolic pressure-pulse interval relationships with an auto threshold at 3 beats in delay. For each mouse, a total of 72 hourly baroreflex sensitivity data points were calculated from the 3 consecutive days of BP data and then averaged to generate one 24-h profile. For *db/db*-ATRF group, 2 out of the 144 data points were excluded as they are identified as outliers by an outlier calculator (https://www.miniwebtool.com/outlier-calculator/).

### Assessment of the prazosin-induced BP drop

Baseline BP was collected from 18-21 week-old control, *db/db*-ALF or *db/db*-ATRF (2 weeks) mice at ZT 5 or ZT17 for one h. Then an α1-adrenergic receptor antagonist, prazosin, was injected (i.p., 1 mg/kg body weight), and BP data was collected for an additional 2 hours. The average BP in the one h prior to prazosin injection and the lowest BP after prazosin injection was selected to quantify the BP responses to prazosin.

### Quantitative analysis of mRNA expression

The 21-week-old db/db and control mice were fed ATRF or ALF for 4 weeks and then euthanized at ZT5, 11, 17, and 23. The liver, mesenteric arteries (MA), kidney, heart, and adrenal gland were collected in RNAlater solution. The fat surrounding the MA was carefully removed under a microscope. The mRNA levels of various clock genes were quantified by real-time PCR, as previously described (53). The real-time PCR primers for each gene were described in Supplementary Table 1.

### Cosinor analysis of Circadian rhythm

The amplitude, robustness, and acrophase of the circadian rhythms in MAP, SBP, DBP, HR, and locomotor activity were determined by the cosinor analysis software (Circadian Physiology, Second Edition, Dr. Robero Refinetti).

### Statistical analysis

All data were expressed as mean ± SEM. The sample size was described in the figures and figure legends as biological replicates. For comparison of 1 parameter between 2 groups of mice, 2-tail Student’s t-test was used. For comparison of 1 parameter between more than 2 groups of mice, one-way ANOVA with Tukey’s post-test was used. For comparison of two parameters between ≥2 groups of mice, two-way ANOVA with Bonferroni’s or Tukey’s post-test as recommended by the software were performed. For comparison of three parameters between ≥2 groups of mice, three-way ANOVA with Tukey’s post-test was performed. All statistical analysis was performed by Prism 8 software (GraphPad Software, San Diego, CA). P < 0.05 was defined as statistically significant.

## Author Contributions

T.H. and W.S.: investigation, data curation, and methodology; T.H., M.J.D: contributed to the conceptualization, review, and editing; V.A.O: contributed to the statistical analysis; M.C.G. and Z.G.: contributed to the conceptualization, supervision, writing, project administration, and funding acquisition.

## Acknowledgments

The authors thank Dr. Wendy Katz for assistance with the indirect calorimetry measurements.

