## Supplemental Data for "Active Time-Restricted Feeding Protects the Blood Pressure Circadian Rhythm in Diabetic Mice"

Tianfei Hou[1], Wen Su[1], Marilyn J. Duncan[2], Vsevolozhskaya A. Olga[3], Zhenheng Guo[1,5*], and Ming C. Gong[4*]
[1]Departments of Pharmacology and Nutritional Sciences, [2]Neuroscience, [3]Biotatistics, and [4]Physiology, University of Kentucky, Lexington, KY; [5]Research and Development, Lexington Veterans Affairs Medical Center, Lexington, KY, USA

***To whom correspondence should be addressed:**

Ming C. Gong, Ph.D.
509 Wethington Building
900 South Limestone Street
Lexington, KY 40536


Zhenheng Guo, Ph.D.
515 Wethington Building
900 South Limestone Street
Lexington, KY 40536


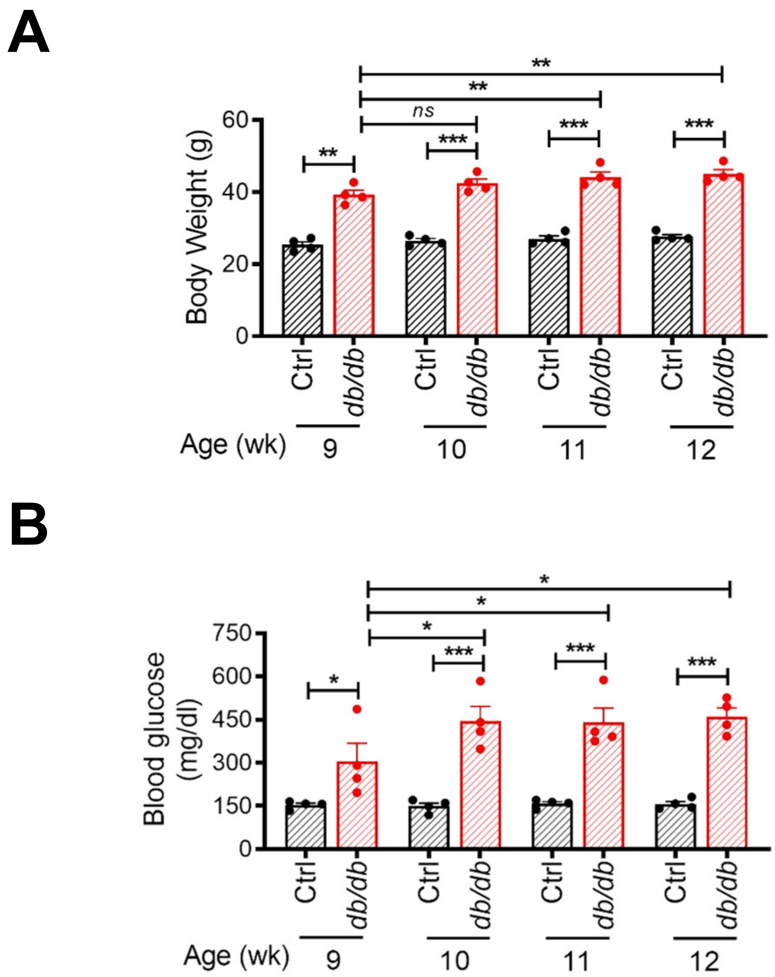


**Supplemental Figure 1. Time course of body weight and nonfasting blood glucose in *db/db* and control mice.** Body weight (**A**) and nonfasting blood glucose (**B**) in 9-, 10-, 11-, and 12-week-old type 2 diabetic *db/db* mice and age-matched nondiabetic control mice (*db/*+). The data were analyzed by 2-way ANOVA with multiple comparisons test. *, p<0.05; **, p<0.01; ***, p<0.001; ns, not significant.

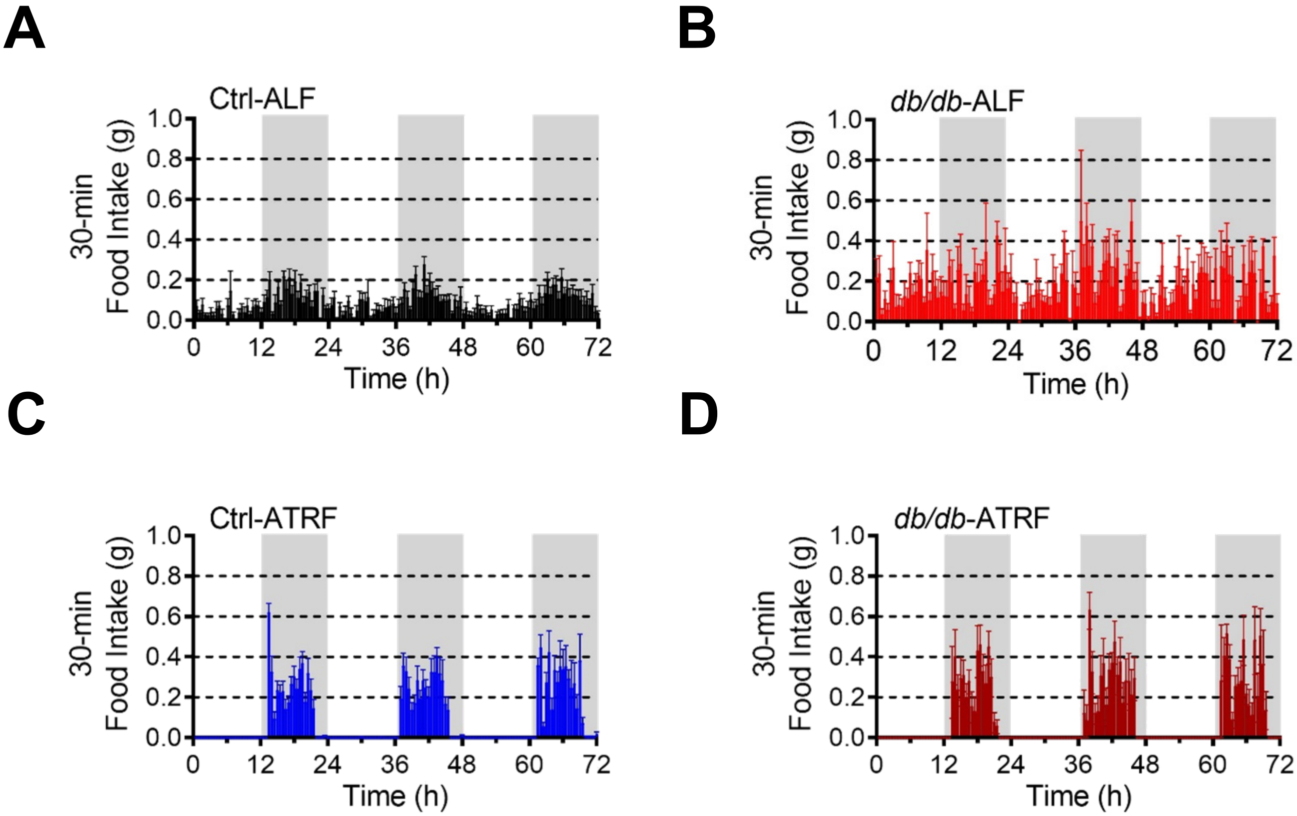


**Supplemental Figure 2. Daily profiles of food intake in the *db/db* and control mice under ALF and ATRF.** Six-week-old *db/db* and control (*db/*+) mice were fed *ad libitum* feeding (ALF) or 8-hour (h) active time-restriction feeding (ATRF). Food intake was monitored by indirect calorimetry in *db/db* and control mice after 10-12 weeks of ATRF or ALF. The data were expressed as the 30-minute intervals in Ctrl-ALF (A; n=10), *db/db*-ALF (B; n=5), Ctrl-ATRF (C; n=10), and *db/db*-ATRF (D; n=5) mice. The grey box indicates the dark-phase.

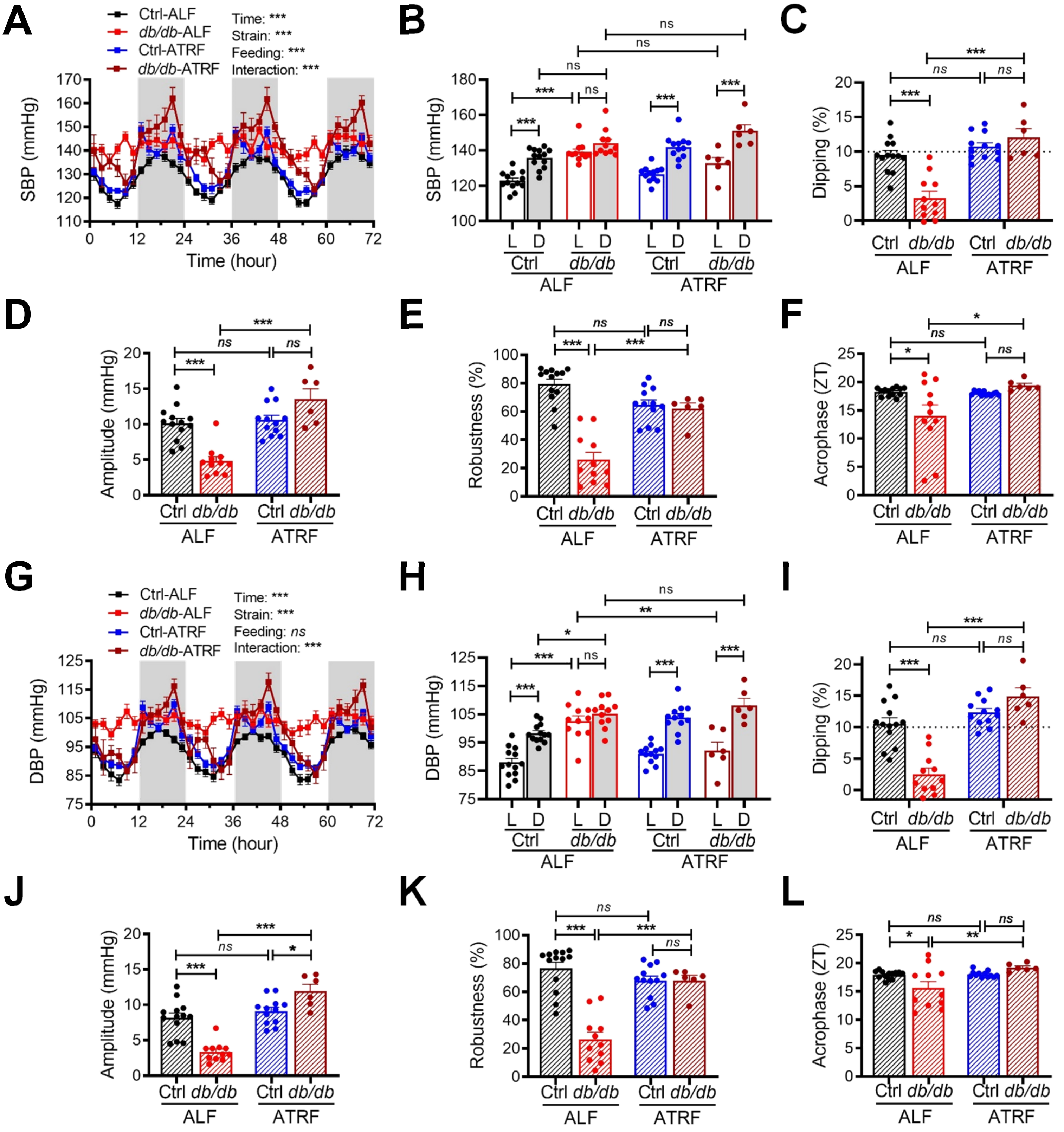


**Supplemental Figure 3. ATRF protects *db/db* mice from systolic blood pressure (SBP) and diastolic blood pressure (DBP) nondipping.** Six-week-old *db/db* and control (*db*/+) mice were fed ALF or 8-h ATRF for 10 weeks. (**A** and **G**) Daily profiles of SBP (A) and DBP (G) in 2-h intervals over 72 hours in Ctrl-ALF (n=13), Ctrl-ATRF (n=12), *db/db*-ALF (n=11), and *db/db*-ATRF (n=6) mice. The grey box indicates the dark phase. (**B** and **H**) The 12-h average SBP (B) and DBP (H) during the light (L) and dark (D) phase. (**C** and **I**) SBP (C) and DBP (I) dipping. The dashed line indicates 10% dipping. (**D** and **J**) Amplitude of the SBP (D) and DBP (J) circadian rhythms. (**E** and **K**) Robustness of the SBP (E) and DBP (K) circadian rhythms. (**F** and **L**) Acrophase of the SBP (F) and DBP (L) circadian rhythms. ZT, zeitgeber time (ZT0 = lights-on and ZT12 = lights-off). The data were analyzed by 3-way ANOVA (A, B, G, and H) and 2-way ANOVA (C–F and I–L) with multiple comparisons test. *, p<0.05; **, p<0.01; ***, p<0.001; ns, not significant.

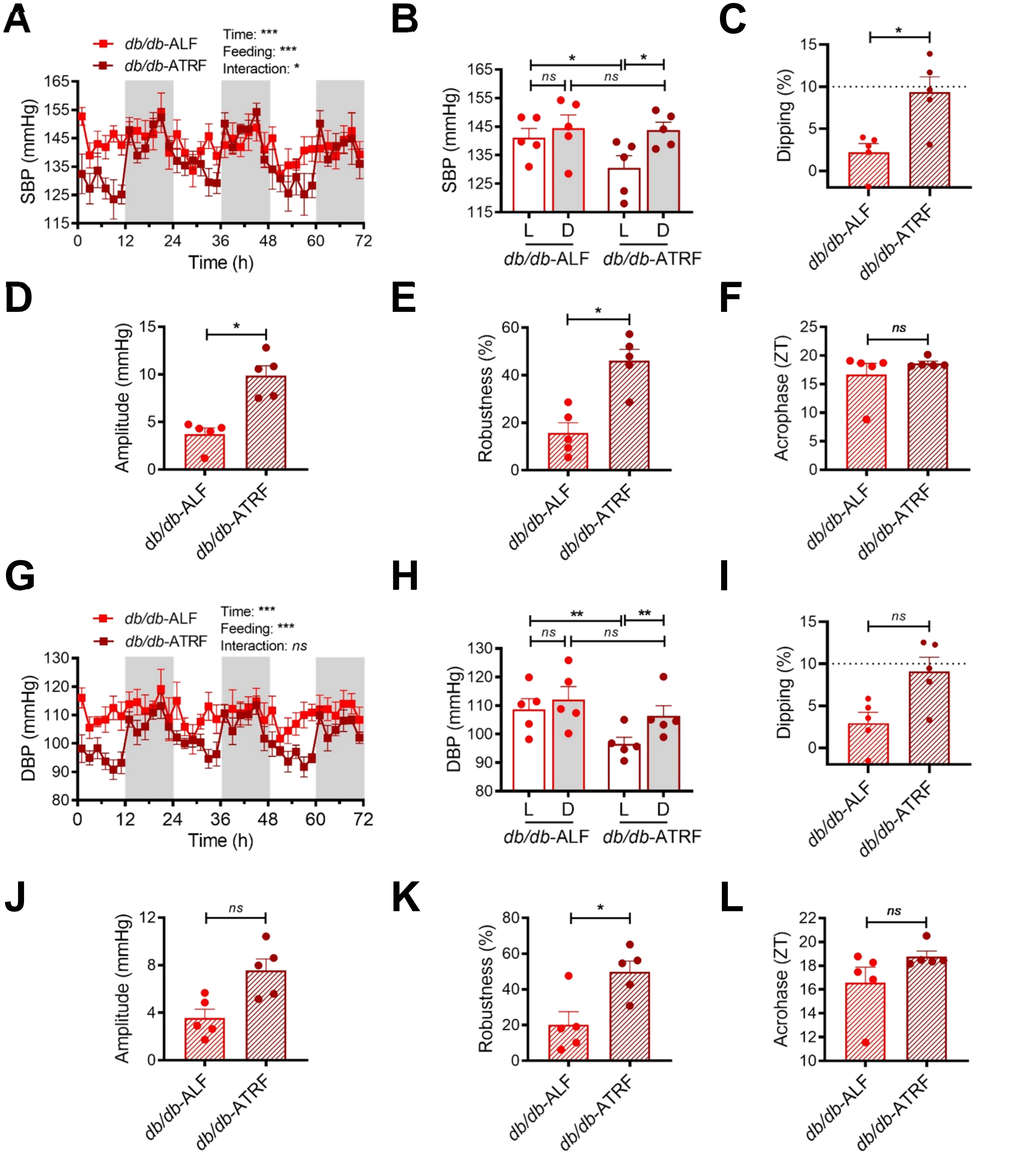


**Supplemental Figure 4. ATRF restores systolic blood pressure (SBP) and diastolic blood pressure (DBP) dipping in *db/db* mice.** SBP and DBP data were first collected in 15-week-old *db/db* mice under ALF and then recorded 9 days after ATRF. Five mice in each group. (**A** and **G**) Daily profiles of SBP (A) and DBP (G) in 2-h intervals in *db/db*-ALF and *db/db*-ATRF mice. The grey box indicates the dark-phase. (**B** and **H**) The 12-hour average SBP (B) and DBP (H) during the light (L) and dark (D) phase. (**C** and **I**) SBP (C) and DBP (I) dipping. The dashed line indicates 10% dipping. (**D** and **J**) Amplitude of the SBP (D) and DBP (J) circadian rhythms. (**E** and **K**) Robustness of the SBP (E) and DBP (K) circadian rhythms. (**F** and **L**) Acrophase of the SBP (F) and DBP (L) circadian rhythms. ZT, zeitgeber time (ZT0 = lights-on and ZT12 = lights-off). The data were analyzed by 2-way ANOVA (A, B, G, and H) with multiple comparisons test or paired t-test (C–F and I–L). *, p<0.05; **, p<0.01; ***, p<0.001; ns, not significant.

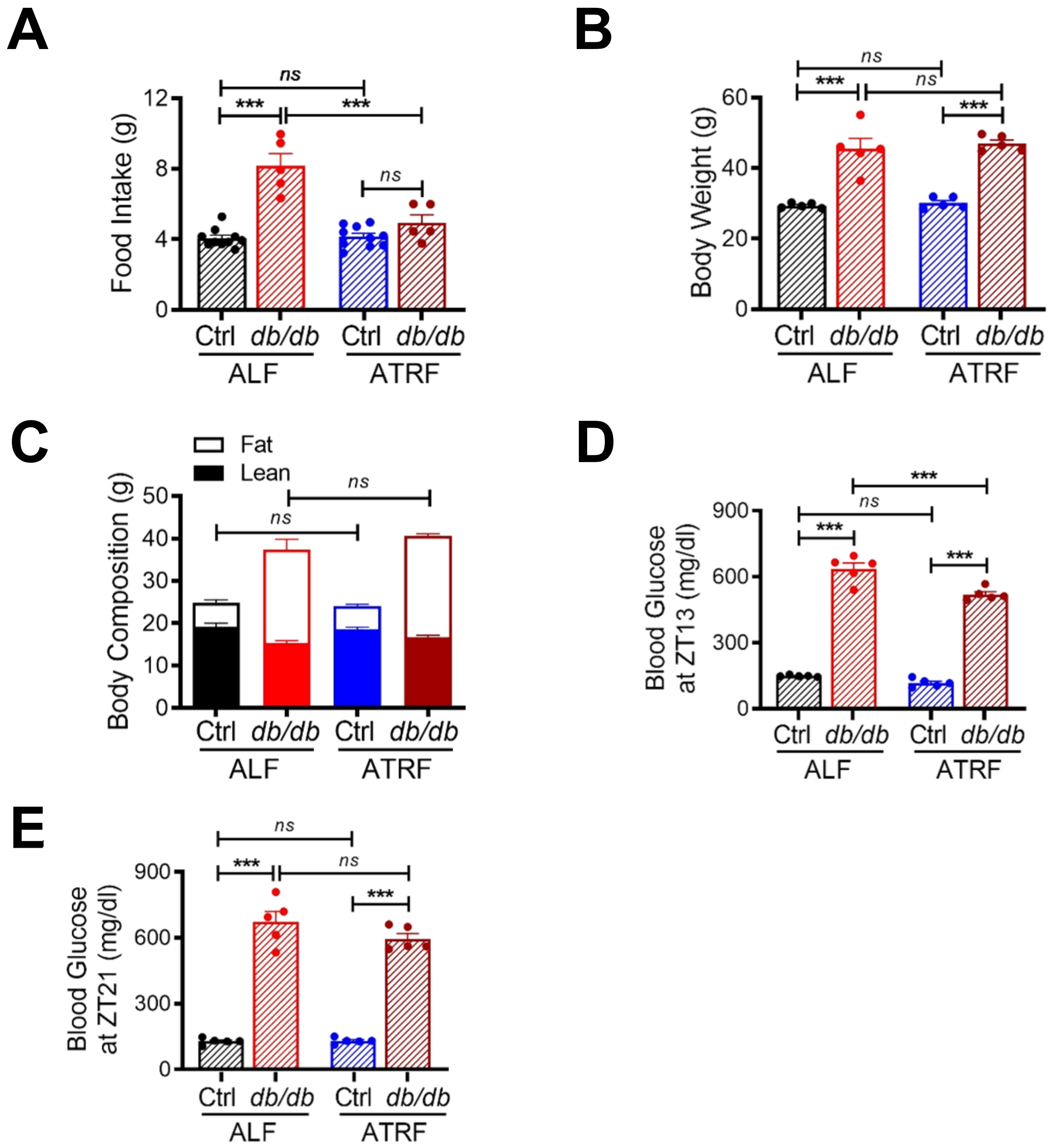


**Supplemental Figure 5. Effects of ATRF on body weight, body composition, and blood glucose in *db/db* and control mice.** Six-week-old *db/db* and control (*db/+*) mice were fed ALF or 8-h ATRF. (**A**) Daily food intake was measured after 10-12 weeks of ALF or 8-h ATRF in Ctrl-ALF (n=10), *db/db*-ALF (n=5), Ctrl-ATRF (n=10) and *db/db*-ATRF (n=5) mice. (**B**–**E**) Body weight (B), body composition (C), and nonfasting blood glucose at ZT13 (D) and ZT21 (E) were measured after 7-8 weeks of ALF or 8-h ATRF. ZT, zeitgeber time (ZT0 = lights-on and ZT12 = lights-off). Five mice in each group. The data were analyzed by 2-way ANOVA with multiple comparisons test. ***, p<0.001; ns, not significant.

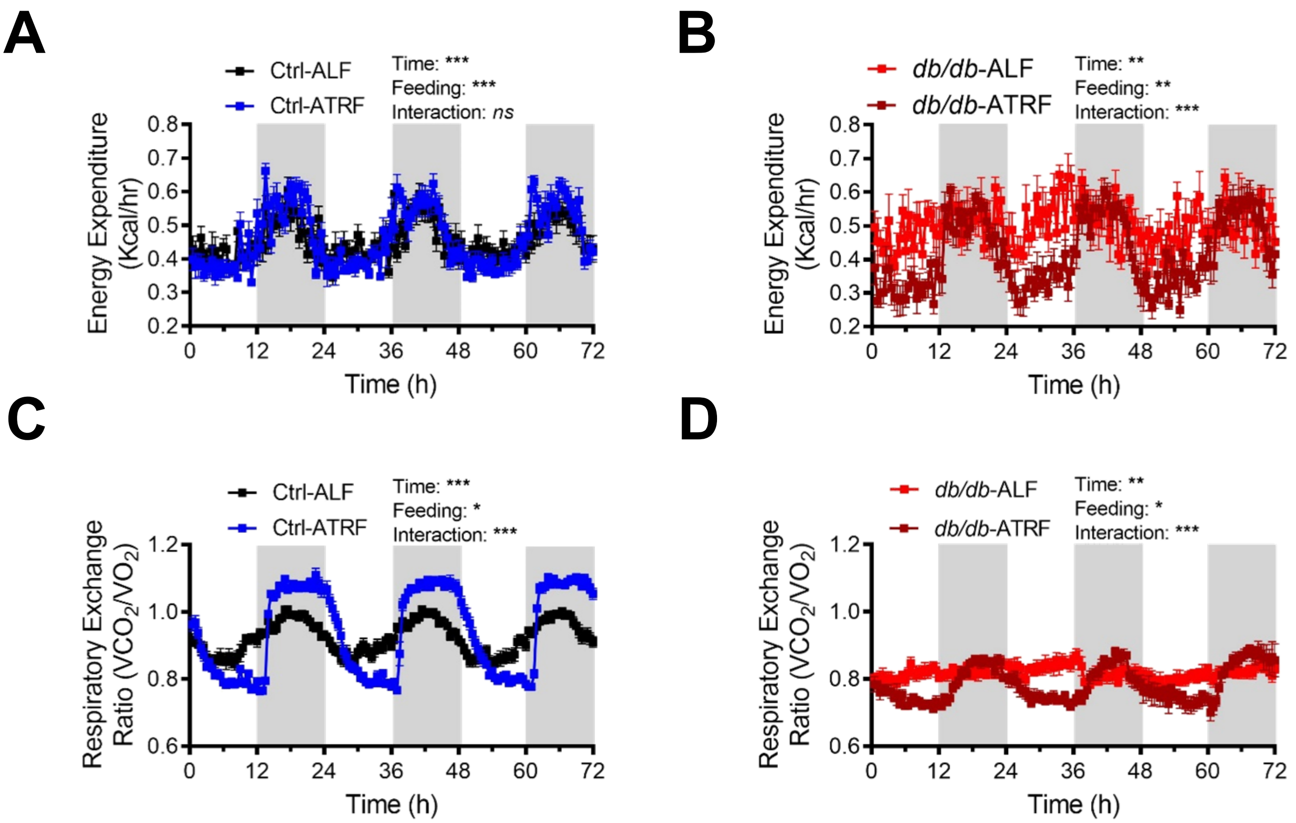


**Supplemental Figure 6. ATRF protects *db/db* mice from the loss of circadian rhythms of energy expenditure and respiratory exchange ratio.** The energy expenditure (EE) and respiratory exchange ratio (REF) were measured by indirect calorimetry in the 16 to 18-week-old *db/db* and control (*db/*+) mice that had been fed ALF or 8-hour ATRF for 10 to 12 weeks. (**A**) Daily profiles of EE in 0.5-h intervals over 72 hours (h) in Ctrl-ALF mice (n=10) and Ctrl-ATRF mice (n=10). (**B**) Daily profiles of EE 0.5-h intervals over 72 h in *db/db*-ALF mice (n=5) and *db/db*-ATRF mice (n=6). (**C**) Daily profiles of RER in 0.5-h intervals over 72 h in Ctrl-ALF mice (n=10) and Ctrl-ATRF mice (n=10). (**D**) Daily profiles of RER in 0.5-h intervals over 72 h in *db/db*-ALF mice (n=5) and *db/db*-ATRF mice (n=6). The grey box indicates the dark phase. The data were analyzed by 2-way ANOVA with multiple comparisons test. *, $p<0.05$; **, $p<0.01$; ***, $p<0.001$; ns, not significant.

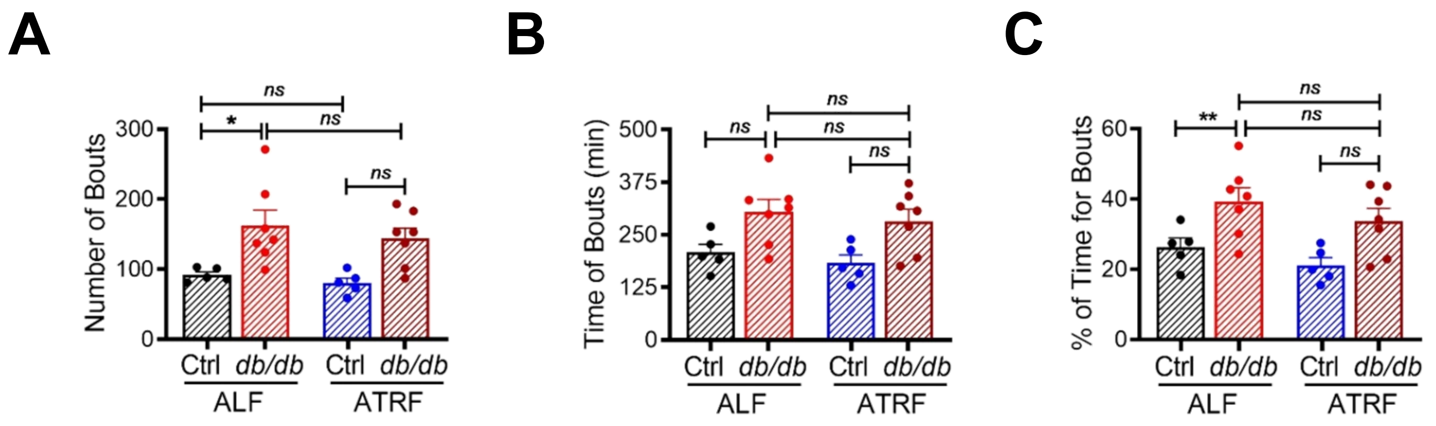


**Supplemental Figure 7. ATRF does not affect the food intake pattern in *db/db* and control mice.** Food intake was monitored by BioDAQ in 14-weeks-old *db/db* and control (*db*/+) mice 4 days after ALF or ATRF. (**A**) Numbers of feeding bouts. (**B**) Duration (time) of feeding bouts. (**C**) Percent of time for feeding bouts. The data were analyzed by 2-way ANOVA with multiple comparisons test. *, p<0.05; **, p<0.01; ns, not significant.

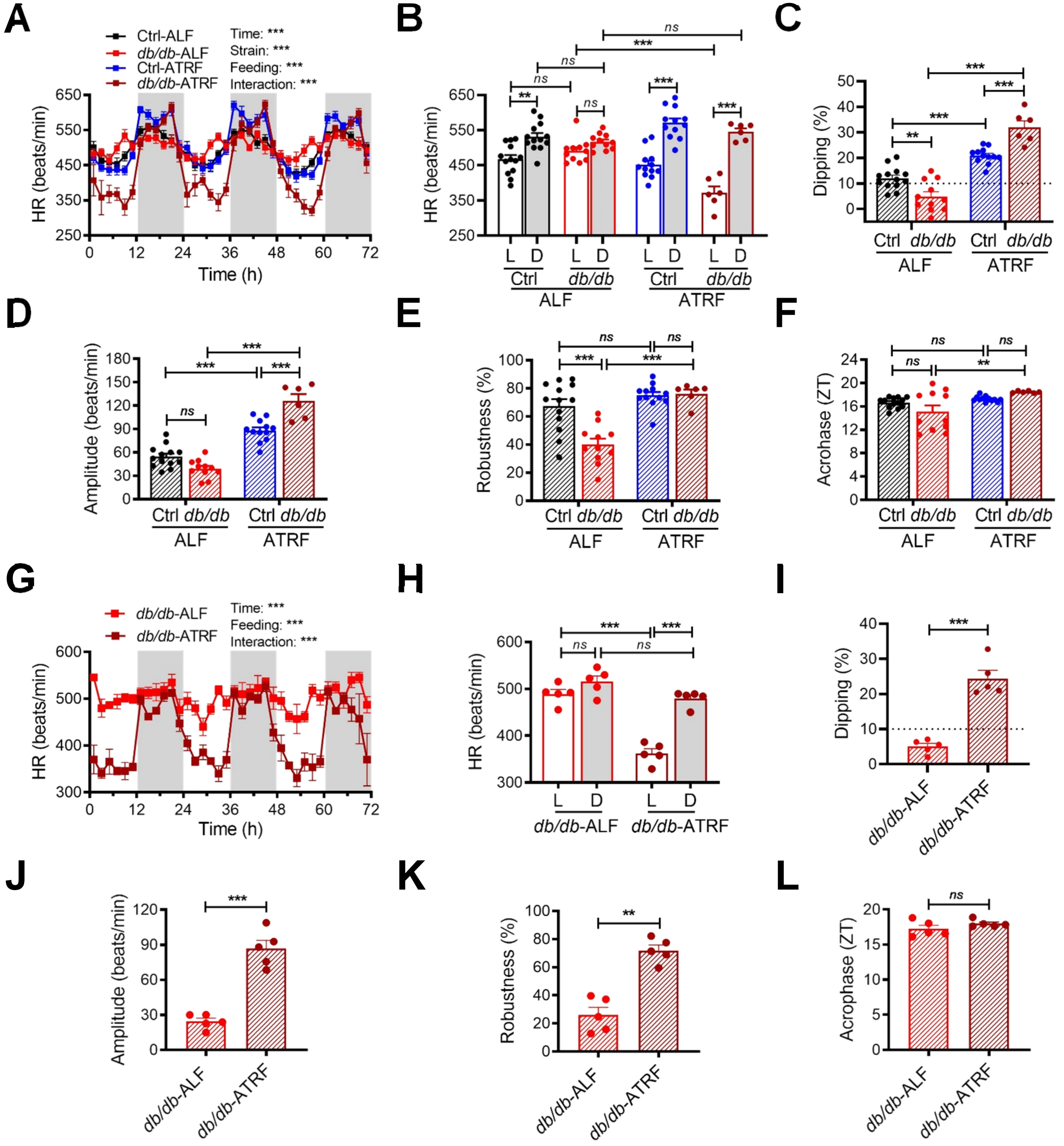


**Supplemental Figure 8. ATRF protects the heart rate circadian rhythm in *db/db* mice.** Heart rate (HR) was recorded by telemetry in the 16-week-old *db/db* and control (*db/+*) mice that had been fed ATRF or ALF for 10 weeks (A–F) as well as the 17-week-old *db/db* mice that were fed first ALF and then 9 days after ATRF (G–L). (**A** and **G**) Daily profiles of HR in 2-h intervals over 72 hours (h). The grey box indicates the dark phase. (**B** and **H**) The 12-h average HR during the light (L) and dark (D) phase. (**C** and **I**) HR dipping (% of HR decrease during the light phase compared to the dark phase). The dashed line indicates 10% dipping. (**D** and **J**) Amplitude of the HR circadian rhythm. (**E** and **K**) Robustness of the HR circadian rhythm. (**F** and **L**) Acrophase of the HR circadian rhythm. ZT, zeitgeber time (ZT0 = lights-on and ZT12 = lights-off). The data were analyzed by 3-ANOVA (A and B) and 2-way ANOVA (C–H) with multiple comparisons test or paired t-test (I–L). *, p<0.05; **, p<0.01; ***, p<0.001; ns, not significant.

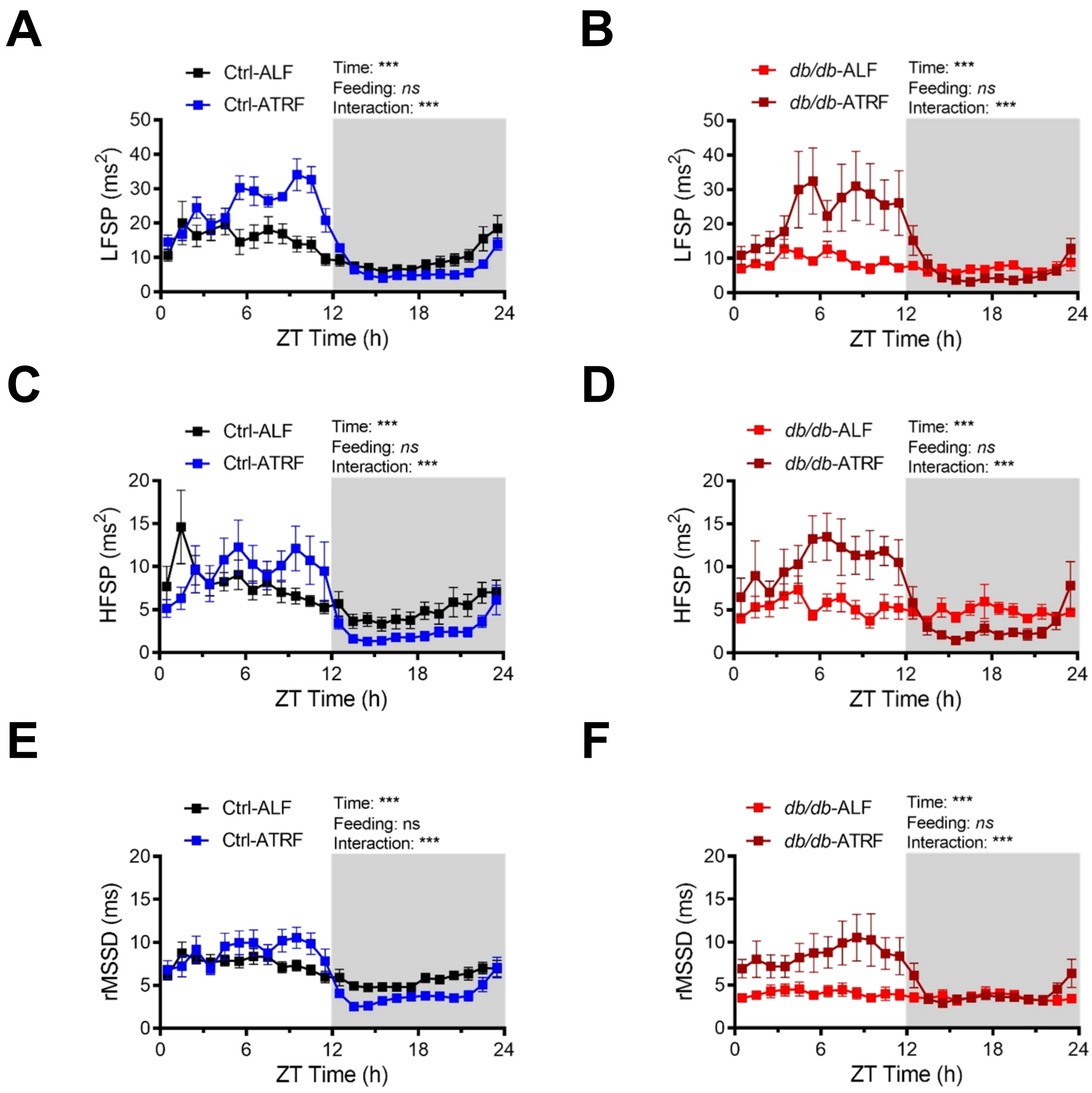


**Supplemental Figure 9. ATRF protects *db/db* mice from the loss of diurnal variation in heart rate variability (HRV).** HRV was analyzed by the low-frequency spectral power (LFSP; A and B), high-frequency spectral power (HFSP; C and D), and root mean square of successive RR interval differences (rMSSD; E and F) in the 16-week-old *db/db* and control mice that had been fed ALF or ATRF for 10 weeks. (**A**) Daily profiles of LFSP in 1-h interval over 24 hours (h) in Ctrl-ALF (n=8) and Ctrl-ATRF (n=8) mice. (**B**) Daily profiles of LFSP in 1-h interval over 24 hours in *db/db*-ALF (n=8) and *db/db*-ATRF (n=6) mice. (**C**) Daily profiles of HFSP in 1-h interval over 24 hours in Ctrl-ALF (n=8) and Ctrl-ATRF (n=8) mice. (**D**) Daily profiles of HFSP in 1-h interval over 24 hours in the *db/db*-ALF (n=8) and *db/db*-ATRF (n=6) mice. (**E**) Daily profiles of rMSSD in 1-h interval over 24 hours in Ctrl-ALF (n=8) and Ctrl-ATRF (n=8) mice. (**F**) Daily profiles of rMSSD in 1-h interval over 24 hours in *db/db*-ALF (n=8) and *db/db*-ATRF (n=6) mice. The grey box indicates the dark phase. ZT, zeitgeber time (ZT0 = lights-on and ZT12 = lights-off). The data were analyzed by 2-way ANOVA. ***, p<0.0001; ns, not significant.

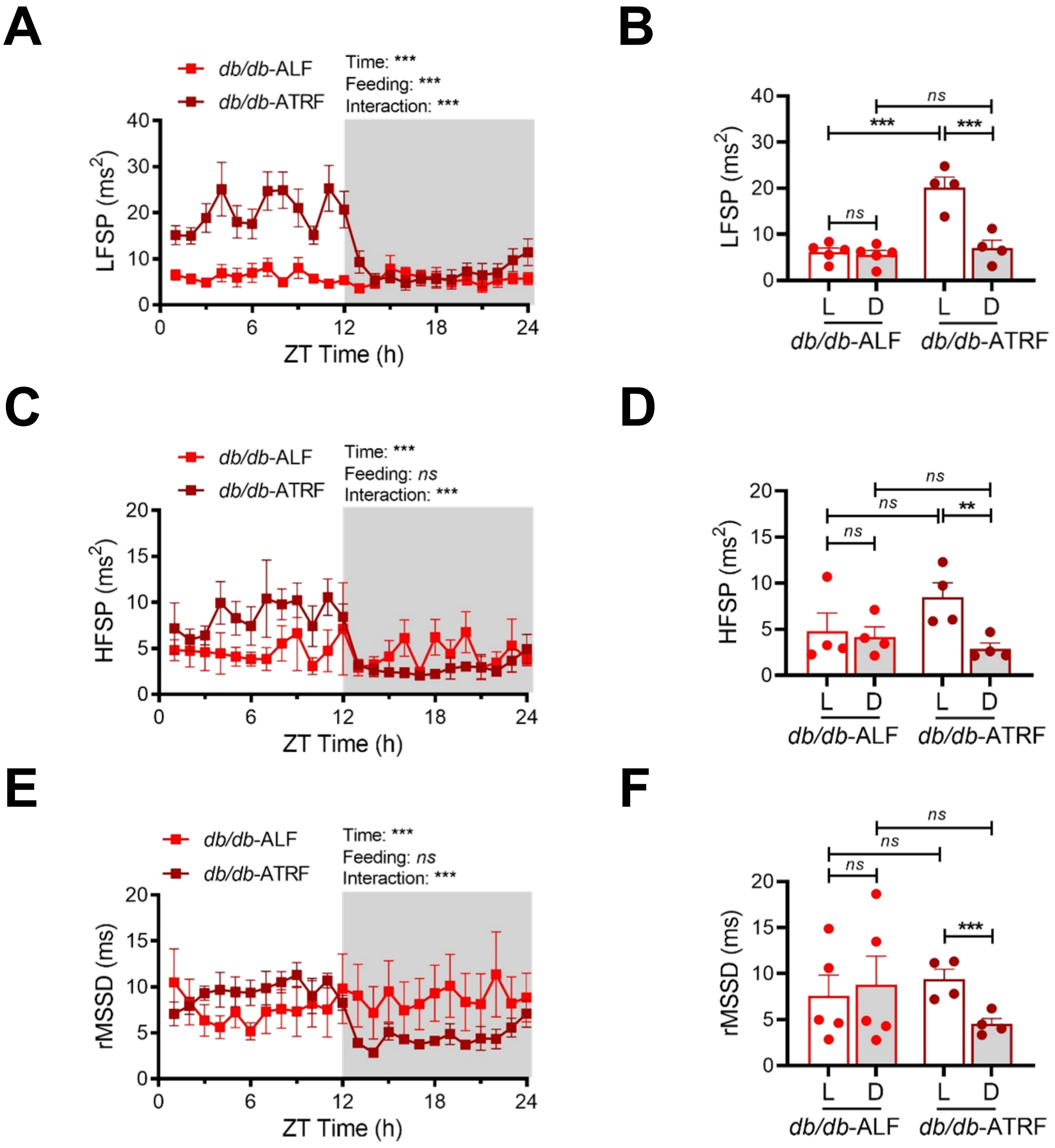


**Supplemental Figure 10. ATRF restores the time-of-day variation of heart rate variability (HRV) in *db/db* mice.** HRV was analyzed by the low-frequency spectral power (LFSP; A and B), high-frequency spectral power (HFSP; C and D), and root mean square of successive RR interval differences (rMSSD; E and F) in the 17-week-old *db/db* mice that had been first fed ALF and then ATRF for 9 days. (**A**) Daily profiles of LFSP in 1-h interval over 24 hours in *db/db*-ALF (n=5) and *db/db*-ATRF (n=4) mice. (**B**) The 12-h average LFSP during the light (L) and dark (D) phase. (**C**) Daily profiles of HFSP in 1-h interval over 24 hours in *db/db*-ALF (n=4) and *db/db*-ATRF (n=4) mice. (**D**) The 12-h average HFSP during the light (L) and dark (D) phase. (**E**) Daily profiles of rMSSD in 1-h interval over 24 hours in *db/db*-ALF (n=5) and *db/db*-ATRF (n=4) mice. (**F**) The 12-h average rMSSD during the light (L) and dark (D) phase. The grey box indicates the dark-phase. ZT, zeitgeber time (ZT0 = lights-on and ZT12 = lights-off). The data were analyzed by 2-way ANOVA with multiple comparisons test. ***, p<0.0001; ns, not significant.  **. P<0.01;  ***, p<0.001; ns, not significant.

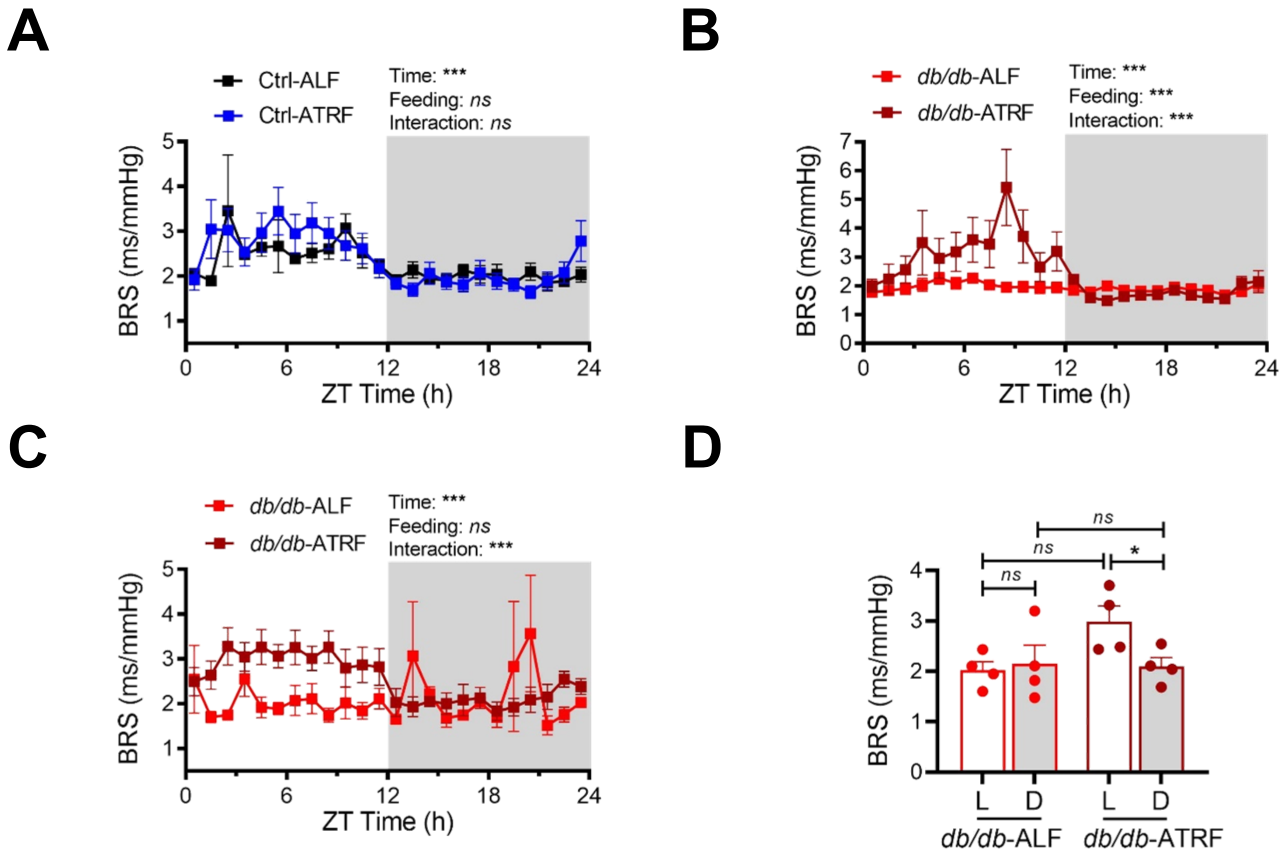


**Supplemental Figure 11. ATRF protects *db/d*b mice from the loss of time-of-day variation of spontaneous baroreflex sensitivity.** Spontaneous baroreflex sensitivity (BRS) was analyzed in the 16-week-old *db/db* and control (*db/*+) mice that had been fed ALF or ATRF for 10 weeks (A and B) or 17-week-old *db/db* mice that had been fed ALF first and then ATRF for 9 days. (**A**) Daily profiles of BRS in 1-h interval over 24 hours in Ctrl-ALF (n=4) and Ctrl-ATRF (n=5) mice. (**B**) Daily profiles of BRS in 1-h interval over 24 hours in *db/db*-ALF (n=10) and *db/db*-ATRF (n=6) mice. (**C**) Daily profiles of BRS in 1-h interval over 24 hours in *db/db*-ALF (n=4) and *db/db*-ATRF (n=4) mice. (**D**) The 12-h average BRS during the light (L) and dark (D) phase in *db/db*-ALF and *db/db*-ATRF mice. The grey box indicates the dark-phase. ZT, zeitgeber time (ZT0 = lights-on and ZT12 = lights-off). The data were analyzed by 2-way ANOVA with multiple comparisons test. *. P<0.05;  ***, p<0.001; ns, not significant.

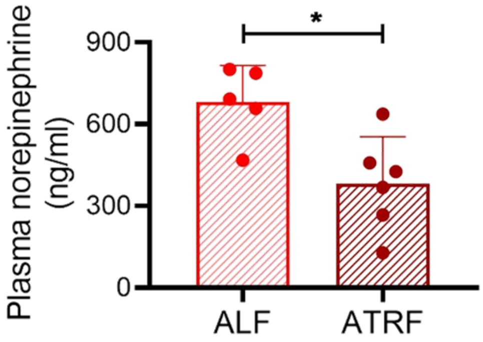


**Supplemental Figure 12. ATRF decreases the plasma norepinephrine (NE) level during the light phase in *db/db* mice.** Plasma was collected at ZT6 during the light phase from the 16-week-old *db/db* mice that had been fed ALF or ATRF for 7 days. ZT, zeitgeber time (ZT0 = lights-on and ZT12 = lights-off). Plasma NE were measured by ELISA. The data was analyzed by unpaired student t-test. *, $p<0.05$.

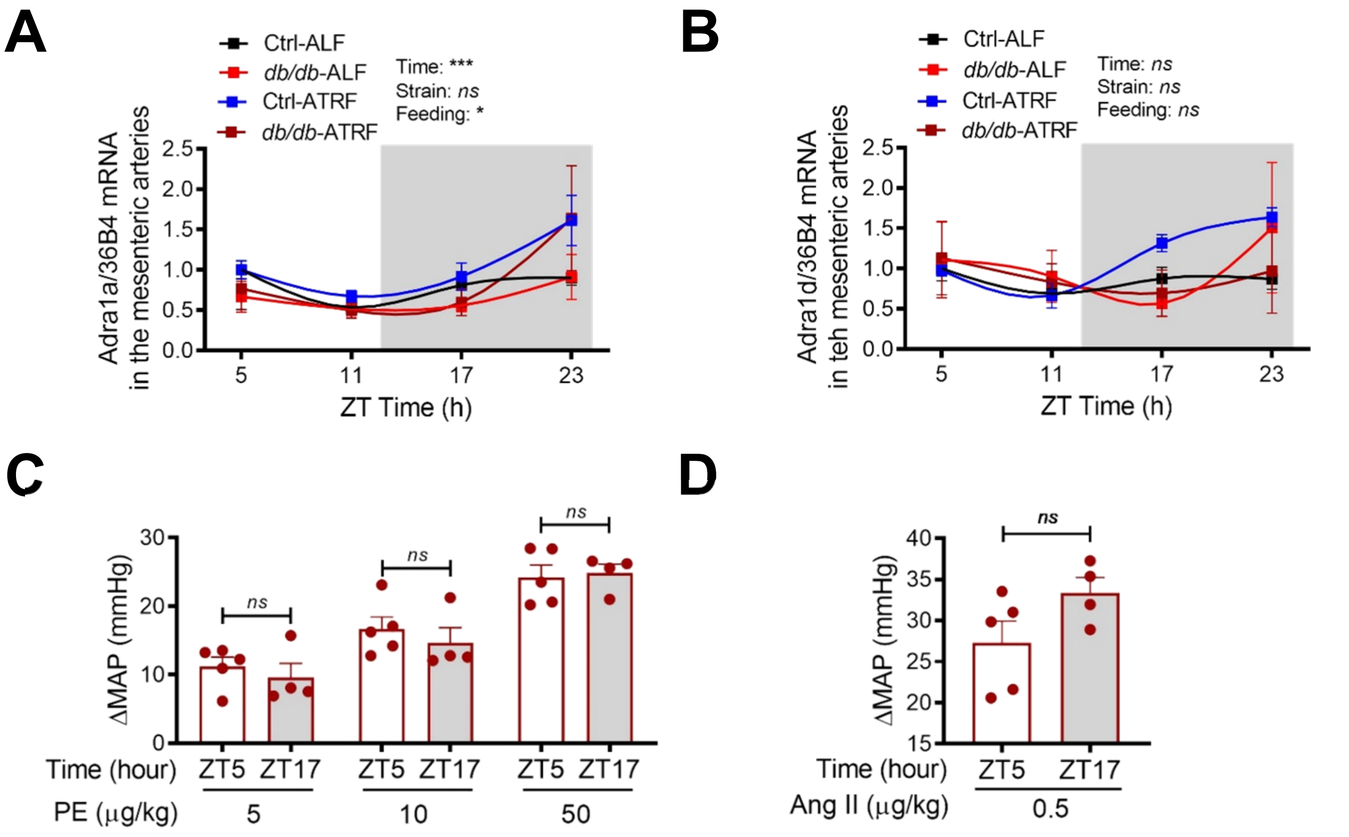


**Supplemental Figure 13. ATRF has no effect on the time-of-day variation of vascular reactivity in *db/db* mice.** (**A** and **B**) The daily mRNA oscillations of adrenergic receptor $\alpha$-1a (Adra1a; A) and adrenergic receptor $\alpha$-1d (Adra1d; B) were determined by real-time PCR in the mesenteric arteries isolated at ZT5, ZT11, ZT17 and ZT23 in 21-week-old *db/db and* control (*db/+*) mice that had been fed ALF or ATRF for 4 weeks. ZT, zeitgeber time (ZT0 = lights-on and ZT12 = lights-off). Adra1a and Adra1d mRNA daily oscillations were normalized to the house-keeping gene 36B4 mRNA and then to the corresponding gene in Ctrl-ALF mice at ZT5. Ctrl-ALF: n=6-7; Ctrl-ATRF: n=4-5; *db/db*-ALF: n=4-5, *db/db*-ATRF: n=3-5. (**C** and **D**) Eighteen-week-old *db/db* mice were first implanted with telemetry and then fed ATRF for 9 days. Under anesthesia conditions, basal MAP was recorded at ZT5 or ZT17 as a baseline. Various concentrations of phenylephrine (PE; C) or a single dose of angiotensin II (Ang II; D) were injected at ZT5 or ZT17 via the femoral artery. The instant pressor responses to PE or Ang II were recorded at ZT5 and ZT17. The differences in MAP ($\Delta$MAP) were calculated by subtracting the basal MAP from PE- or Ang II-induced maximal MAP at ZT5 or ZT17. The data were analyzed by 3-way ANOVA with multiple comparisons test (A and B) and paired t-test (C and D). *, $p<0.05$; ***, $p<0.001$; ns, not significant.

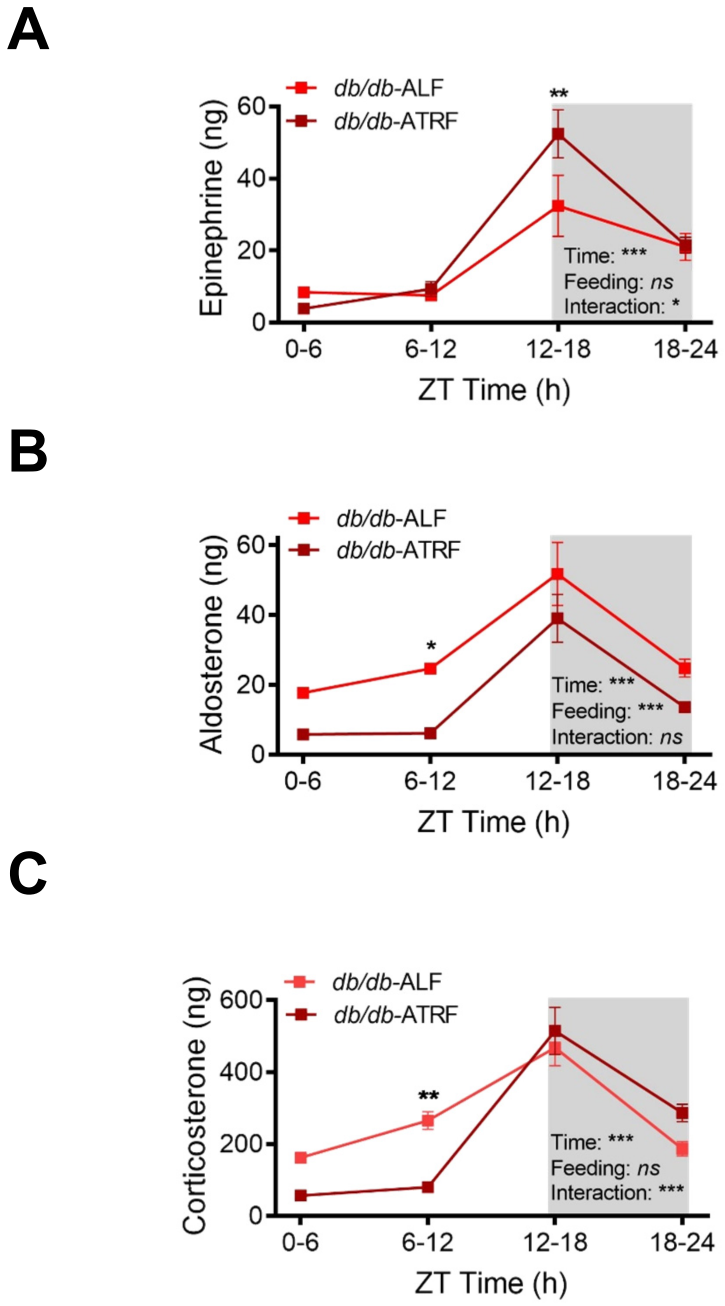


**Supplemental Figure 14. ATRF improves the time-of-day variations in urine epinephrine, aldosterone, and corticosterone in *db/db* mice.** Urines were collected every 6 hours within one 24-hour day at ZT0-6, ZT6-12, ZT12-18, and ZT18-24 from 15-week-old *db/db* mice that had been fed ALF or ATRF for 8 weeks. ZT, zeitgeber time (ZT0 = lights-on and ZT12 = lights-off). The urinary contents of epinephrine (**A**), aldosterone (**B**), and corticosterone (**C**) were measured by ELISA. The data were analyzed by 2-way ANOVA with multiple comparisons test (n=4-5). *, p<0.05; **, p<0.01; ***, p<0.001; ns, not significant.

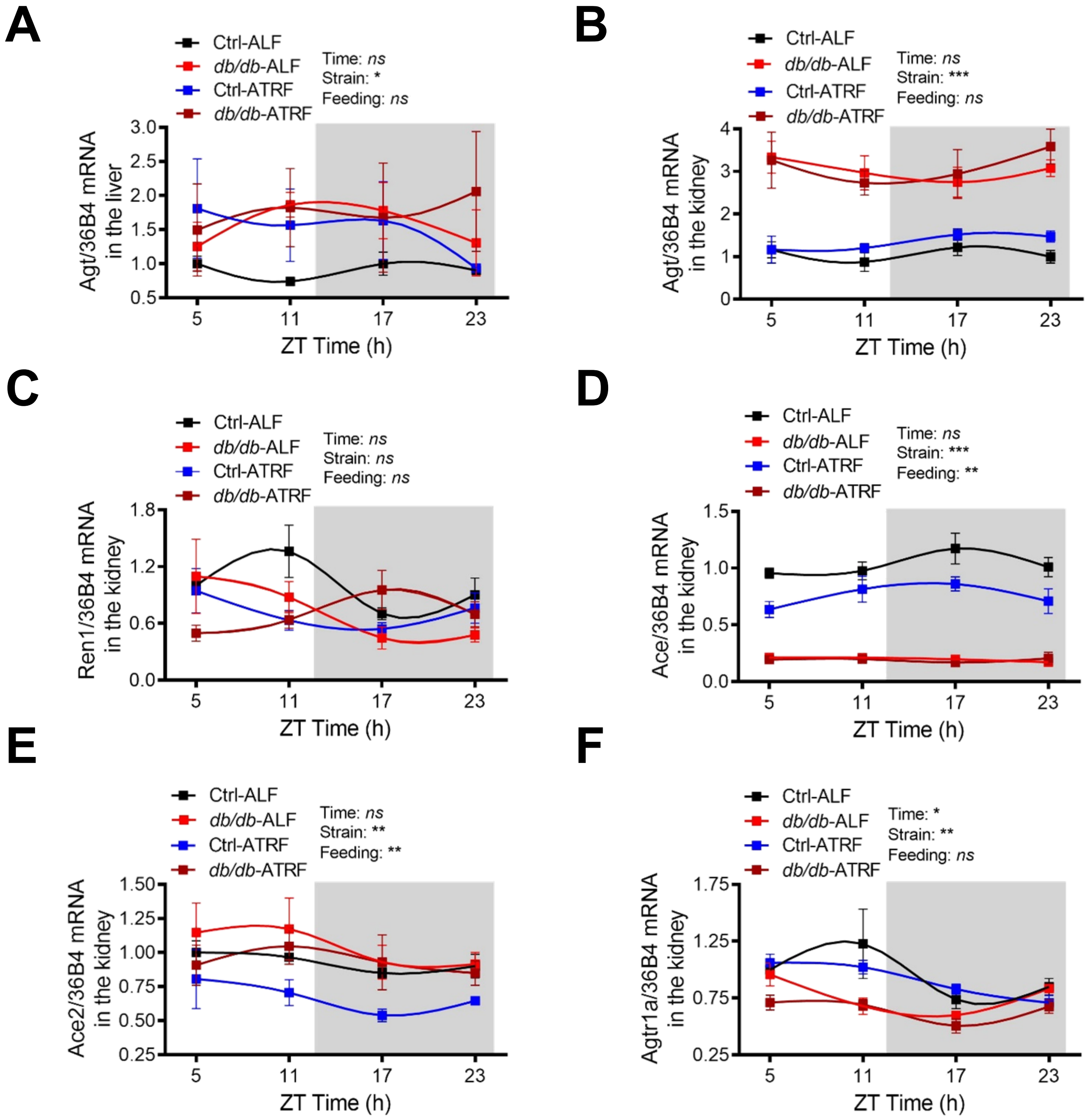


**Supplemental Figure 15. ATRF has no effect on the daily mRNA oscillations of Agt, Ren1, Ace1, Ace2, and Agtr1a in the liver or kidney in *db/db* mice.** The daily mRNA oscillations of angiotensinogen (Agt) in the liver (**A**) and kidney (**B**), renin (Ren1) in the kidney (**C**), angiotensin-converting enzyme (Ace) in the kidney (**D**), Ace2 in the kidney (**E**), and angiotensin II receptor 1a (Agtr1a) in the kidney (**F**) were determined by real-time PCR at zeitgeber time (ZT)5, ZT11, ZT17 and ZT23 in 21-week-old *db/db and* control (*db/+*) mice that had been fed ALF or ATRF for 4 weeks. The expression of each gene was normalized to the house-keeping gene 36B4 mRNA in the same sample and then normalized to the corresponding gene in Ctrl-ALF mice at ZT5. Ctrl-ALF: n=6-7; Ctrl-ATRF: n=4-5; *db/db*-ALF: n=4-5, *db/db*-ATRF: n=3-5. The data were analyzed by 3-way ANOVA with multiple comparisons test. *, p<0.05; **, p<0.01; ***, p<0.001; ns, not significant.

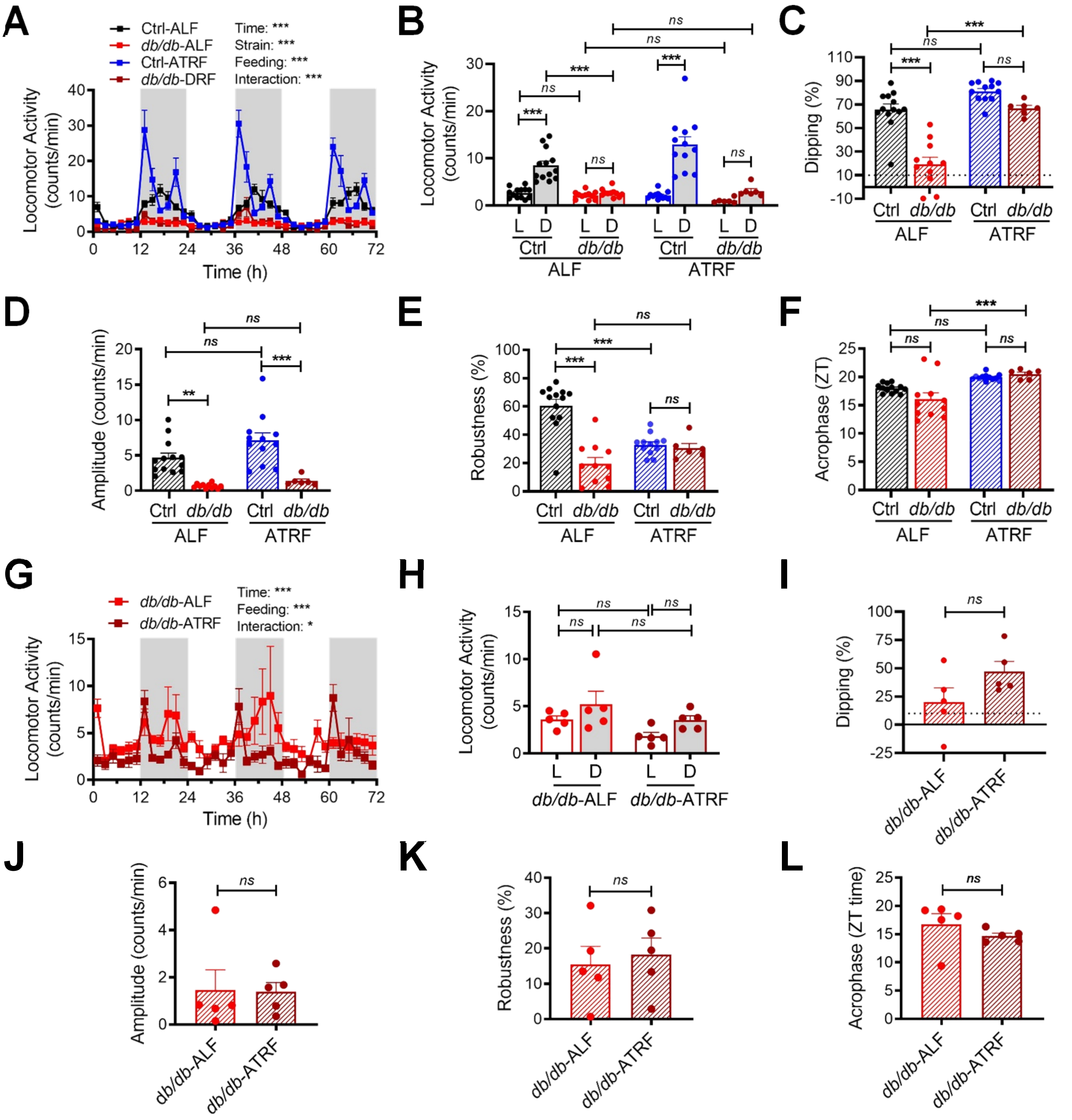


**Supplemental Figure 16. ATRF has no effect on locomotor activity circadian rhythm in *db/db* mice.** Locomotor activity was recorded by telemetry in the 16-week-old Ctrl-ALF (n=13), *db/db*-ALF (n=11), Ctrl-ATRF (n=12), and *db/db*-ATRF (n=6) mice that had been fed ALF or ATRF for 10 weeks (A–F) or 15-weeks-old *db/db* mice that had been fed ALF or ATRF for 9 days. (**A** and **G**) Daily profiles of the locomotor activity in 2-h intervals over 72 hours (h). The grey box indicates the dark phase. (**B** and **H**) The 12-h average locomotor activity during the light (L) and dark (D) phase. (**C** and **I**) Locomotor activity dipping (% of locomotor activity decrease during the light phase compared to the dark phase). The dashed line indicates 10% dipping. (**D** and **J**) Amplitude of the locomotor activity circadian rhythm. (**E** and **K**) Robustness of the locomotor activity circadian rhythm. (**F** and **L**) Acrophase of the locomotor activity circadian rhythm. ZT, zeitgeber time (ZT0 = lights-on and ZT12 = lights-off). The data were analyzed by 3-way ANOVA (A and B) and 2-way ANOVA (C–H) with multiple comparisons test or paired t-test (I–L). *, p<0.05; **, p<0.01; ***, p<0.001; ns, not significant.

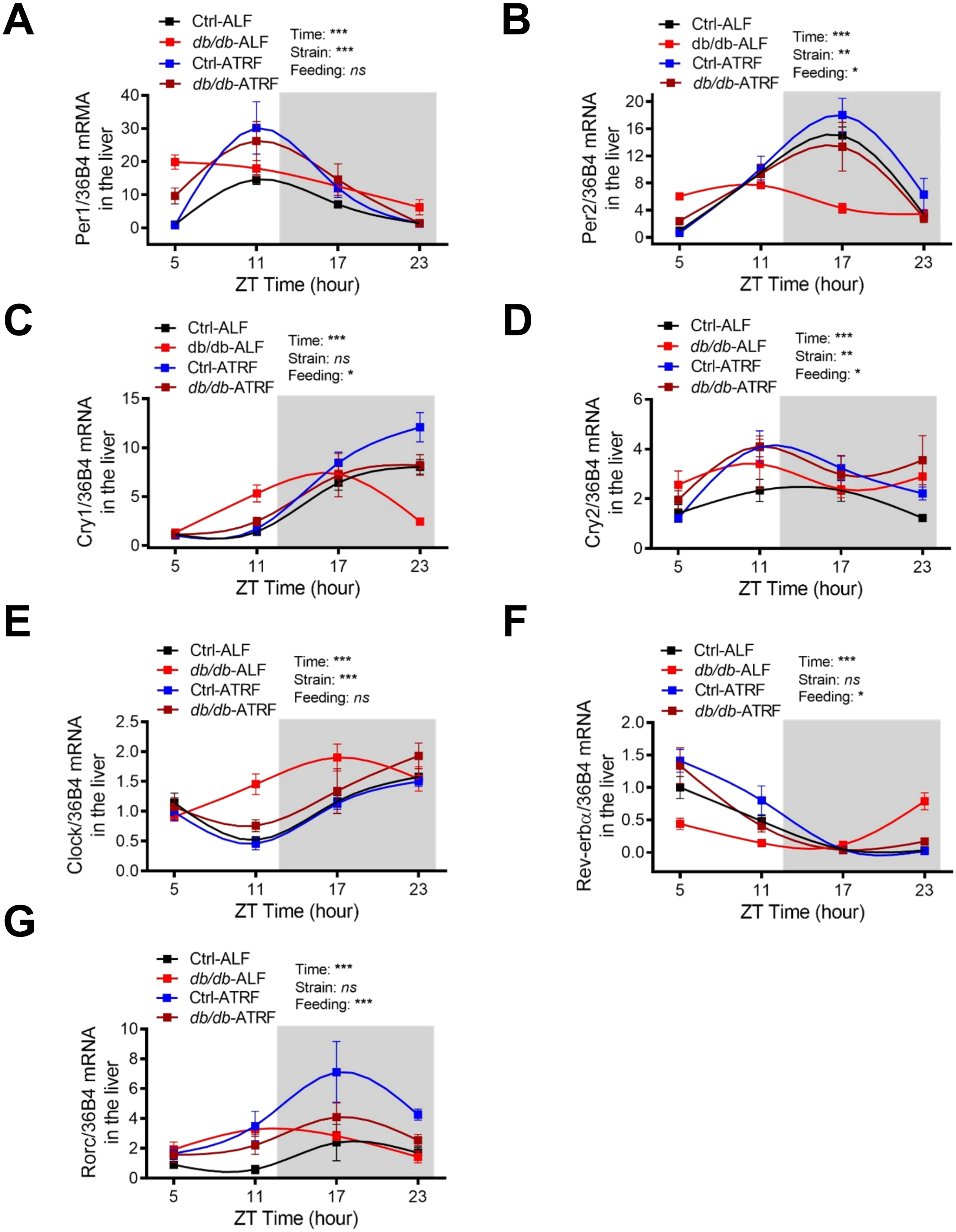


**Supplemental Figure 17. Effects of ATRF on the daily oscillations of clock genes in the liver of *db/db* mice.** The daily mRNA oscillations of period 1 (Per1; **A**), Per2 (**B**), cryptochrome 1 (Cry1; **C**), Cry2 (**D**), Clock (**E**), nuclear receptor subfamily 1, group D, member 1 (Nr1d1; also known as Rev-erbα; **F**), and RAR-related orphan receptor gamma (Rorc; **G**) were determined by real-time PCR in the liver isolated at ZT5, ZT11, ZT17 and ZT23 from 21-week-old *db/db* and control (*db/+*) mice that had been fed ALF or ATRF for 4 weeks. ZT, zeitgeber time (ZT0 = lights-on and ZT12 = lights-off). The grey box indicates the dark phase. The daily mRNA oscillations of each gene were normalized to the house-keeping gene 36B4 mRNA and then to the corresponding gene in the Ctrl-ALF mice at ZT5. Ctrl-ALF: n=6-7; Ctrl-ATRF: n=4-5; *db/db*-ALF: n=4-5, *db/db*-ATRF: n=3-5. The data were analyzed by 3-way ANOVA with multiple comparisons test. *, $p<0.05$; **, $p<0.01$; ***, $p<0.001$; ns, not significant.

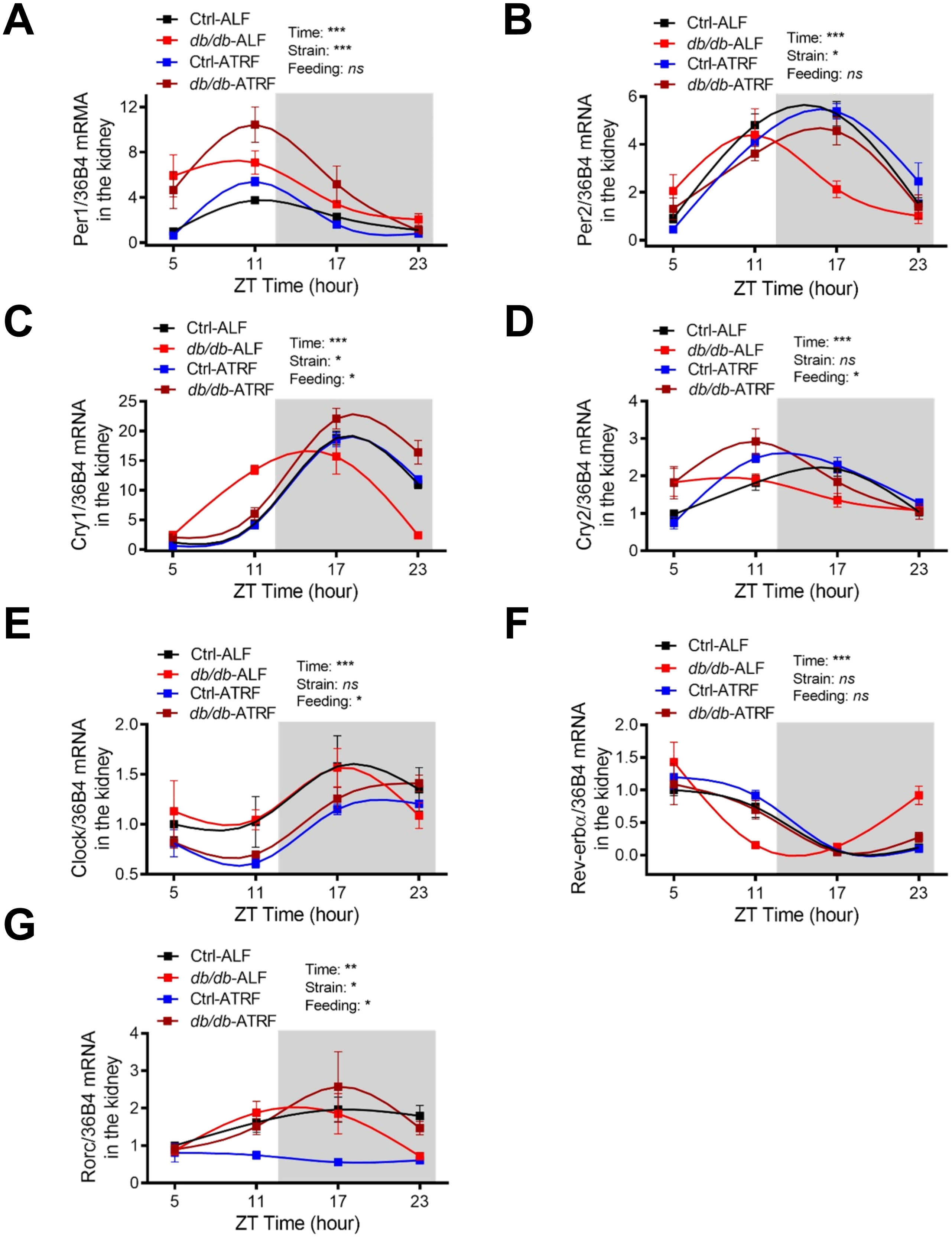


**Supplemental Figure 18. Effects of ATRF on the daily oscillations of clock genes in the kidney in *db/db* mice.** The daily mRNA oscillations of period 1 (Per1; **A**), Per2 (**B**), cryptochrome 1 (Cry1; **C**), Cry2 (**D**), Clock (**E**), nuclear receptor subfamily 1, group D, member 1 (Nr1d1; also known as Rev-erbα; **F**), and RAR-related orphan receptor gamma (Rorc; **G**) were determined by real-time PCR in the kidney isolated at ZT5, ZT11, ZT17 and ZT23 from 21-week-old *db/db* and control (*db/+*) mice that had been fed ALF or ATRF for 4 weeks. ZT, zeitgeber time (ZT0 = lights-on and ZT12 = lights-off). The grey box indicates the dark phase. The daily mRNA oscillations of each gene were normalized to the house-keeping gene 36B4 mRNA and then to the corresponding gene in the Ctrl-ALF mice at ZT5. Ctrl-ALF: n=6-7; Ctrl-ATRF: n=4-5; *db/db*-ALF: n=4-5, *db/db*-ATRF: n=3-5. The data were analyzed by 3-way ANOVA with multiple comparisons test. *, $p<0.05$; **, $p<0.01$; ***, $p<0.001$; ns, not significant.

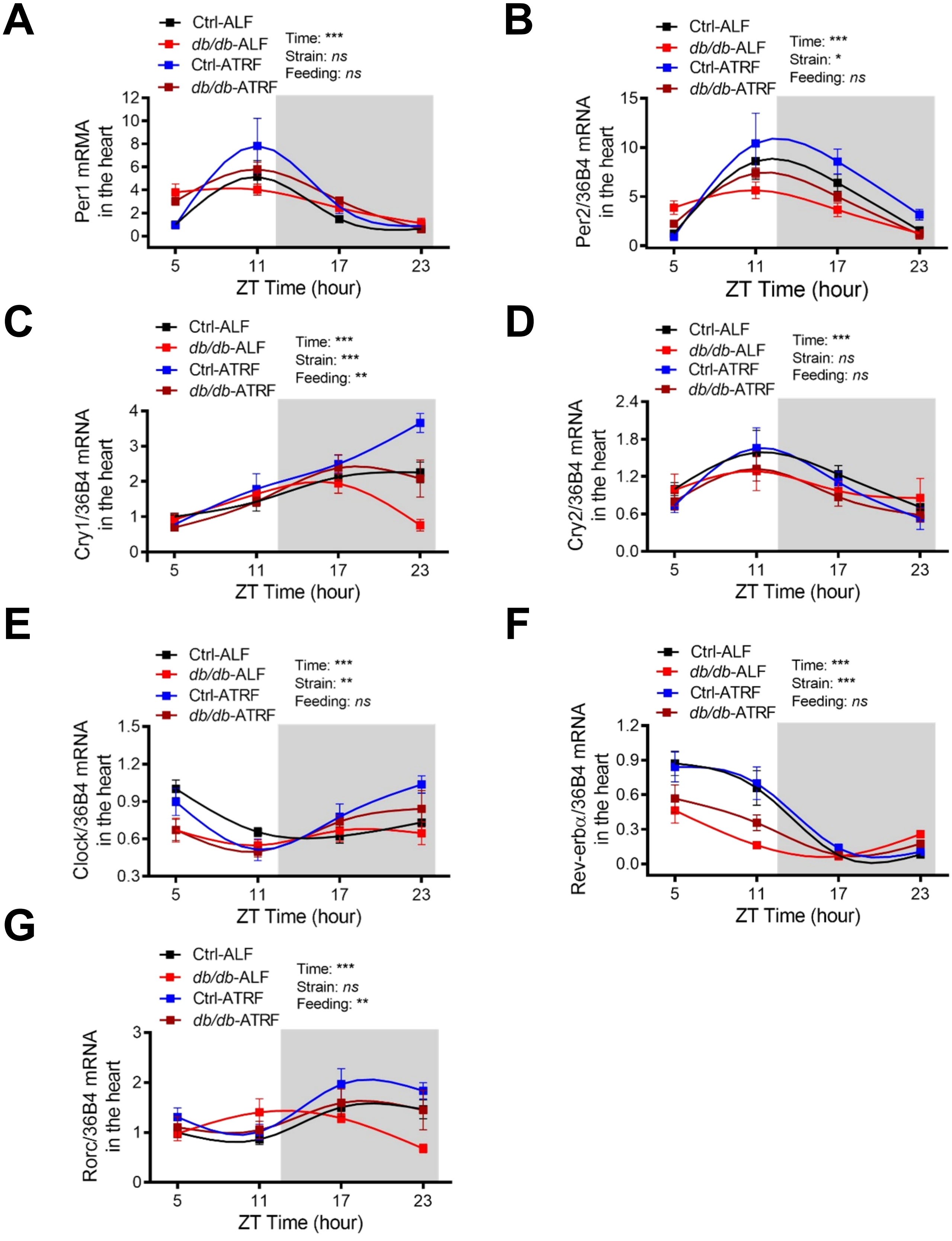


**Supplemental Figure 19. Effects of ATRF on the daily oscillation of clock genes in the heart in *db/db* mice.** The daily mRNA oscillations of period 1 (Per1; **A**), Per2 (**B**), cryptochrome 1 (Cry1; **C**), Cry2 (**D**), Clock (**E**), nuclear receptor subfamily 1, group D, member 1 (Nr1d1; also known as Rev-erbα; **F**), and RAR-related orphan receptor gamma (Rorc; **G**) were determined by real-time PCR in the heart isolated at ZT5, ZT11, ZT17 and ZT23 from 21-week-old *db/db* and control (*db/+*) mice that had been fed ALF or ATRF for 4 weeks. ZT, zeitgeber time (ZT0 = lights-on and ZT12 = lights-off). The grey box indicates the dark phase. The daily mRNA oscillations of each gene were normalized to the house-keeping gene 36B4 mRNA and then to the corresponding gene in the Ctrl-ALF mice at ZT5. Ctrl-ALF: n=6-7; Ctrl-ATRF: n=4-5; *db/db*-ALF: n=4-5, *db/db*-ATRF: n=3-5. The data were analyzed by 3-way ANOVA with multiple comparisons test. *, p<0.05; **, p<0.01; ***, p<0.001; ns, not significant.

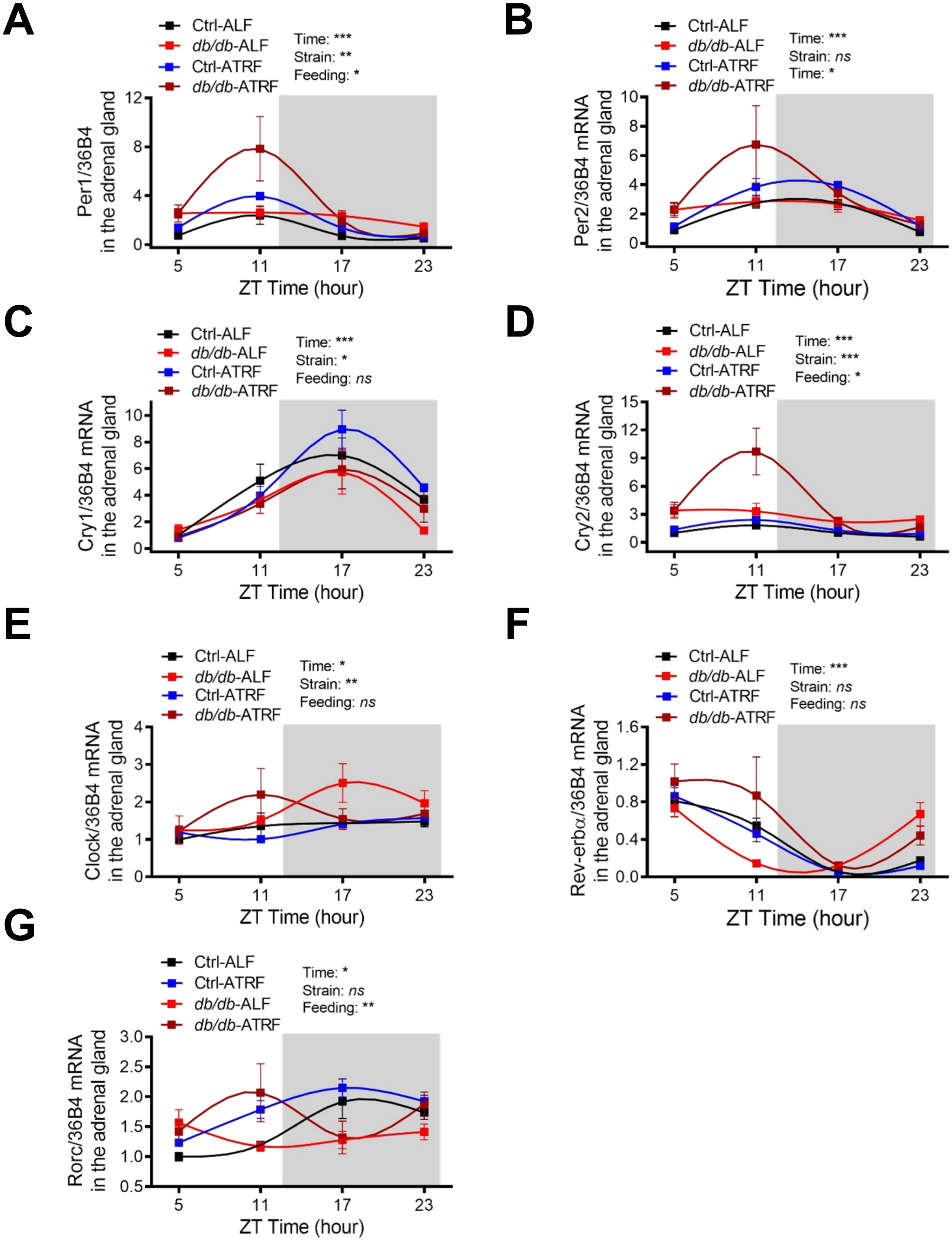


**Supplemental Figure 20. Effects of ATRF on the daily oscillation of clock genes in the adrenal gland in *db/db* mice.** The daily mRNA oscillations of period 1 (Per1; **A**), Per2 (**B**), cryptochrome 1 (Cry1; **C**), Cry2 (**D**), Clock (**E**), nuclear receptor subfamily 1, group D, member 1 (Nr1d1; also known as Rev-erbα; **F**), and RAR-related orphan receptor gamma (Rorc; **G**) were determined by real-time PCR in the adrenal gland isolated at ZT5, ZT11, ZT17 and ZT23 from 21-week-old *db/db* and control (*db/+*) mice that had been fed ALF or ATRF for 4 weeks. ZT, zeitgeber time (ZT0 = lights-on and ZT12 = lights-off). The grey box indicates the dark phase. The daily mRNA oscillations of each gene were normalized to the house-keeping gene 36B4 mRNA and then to the corresponding gene in the Ctrl-ALF mice at ZT5. Ctrl-ALF: n=6-7; Ctrl-ATRF: n=4-5; *db/db*-ALF: n=4-5, *db/db*-ATRF: n=3-5. The data were analyzed by 3-way ANOVA with multiple comparisons test. *, p<0.05; **, p<0.01; ***, p<0.001; ns, not significant.

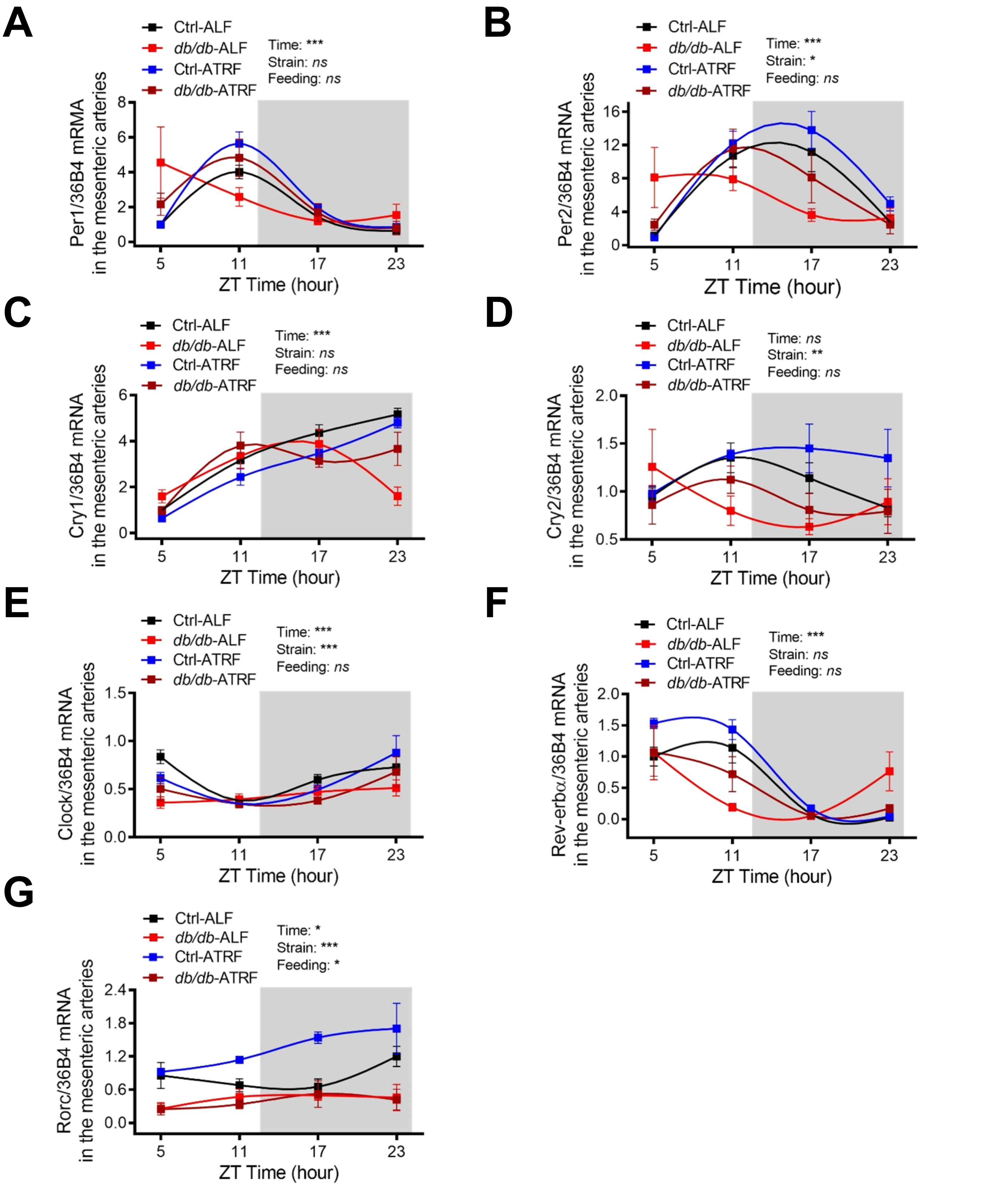


**Supplemental Figure 21. Effects of ATRF on the daily oscillation of clock genes in the mesenteric arteries in *db/db* mice.** The daily mRNA oscillations of period 1 (Per1; **A**), Per2 (**B**), cryptochrome 1 (Cry1; **C**), Cry2 (**D**), Clock (**E**), nuclear receptor subfamily 1, group D, member 1 (Nr1d1; also known as Rev-erb$\alpha$; **F**), and RAR-related orphan receptor gamma (Rorc; **G**) were determined by real-time PCR in the mesenteric arteries isolated at ZT5, ZT11, ZT17, and ZT23 from 21-week-old *db/db* and control (*db/+*) mice that had been fed ALF or ATRF for 4 weeks. ZT, zeitgeber time (ZT0 = lights-on and ZT12 = lights-off). The grey box indicates the dark phase. The daily mRNA oscillations of each gene were normalized to the house-keeping gene 36B4 mRNA and then to the corresponding gene in the Ctrl-ALF mice at ZT5. Ctrl-ALF: n=6-7; Ctrl-ATRF: n=4-5; *db/db*-ALF: n=4-5, *db/db*-ATRF: n=3-5. The data were analyzed by 3-way ANOVA with multiple comparisons test. *, p<0.05; **, p<0.01; ***, p<0.001; ns, not significant.

A

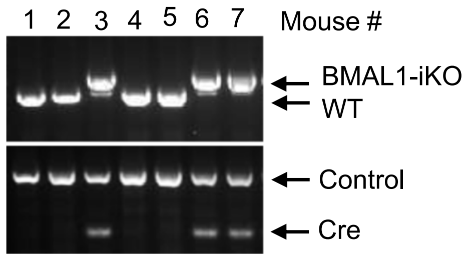


B

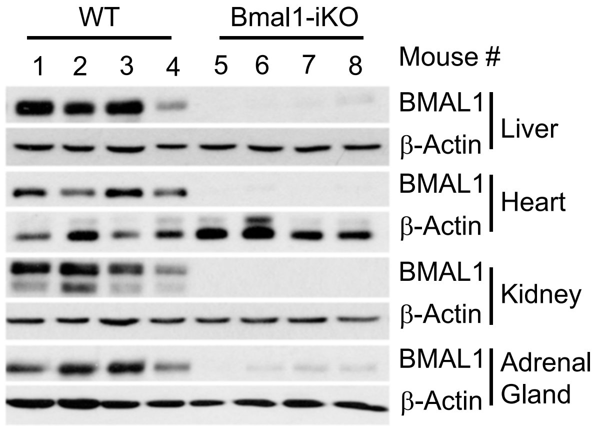


**Supplemental Figure 22. Characterization of the tamoxifen-inducible global Bmal1 knockout mouse model (Bmal1-iKO).** (**A**) Genotyping of Bmal1-iKO mice by genomic PCR from mouse tails. Of mice examined, mouse #1, 2, 4, and 5 were Bmal1 floxed mice, whereas mouse #3, 6, and 7 were Bmal1-iKO. (**B**) Representative Western blots of BMAL1 protein expression in the liver, heart, kidney, and adrenal gland from the Bmal1-iKO and WT ($Bmal1^{fl/fl}$) control mice (n=4).

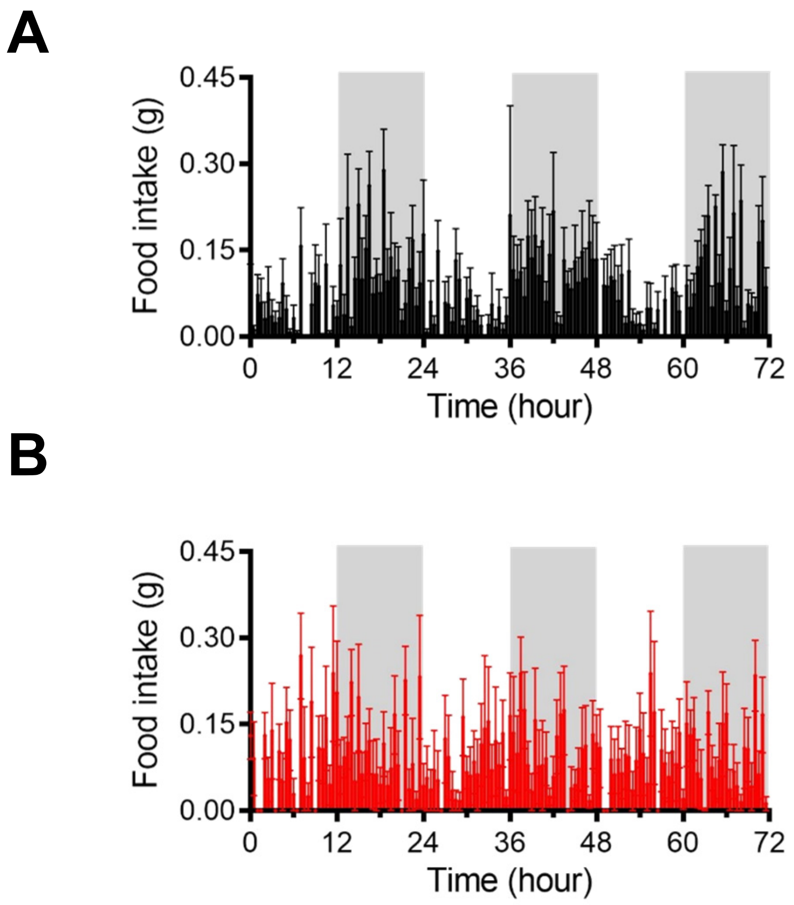


**Supplemental Figure 23. Food intake diurnal rhythm in Bmal1-iKO and WT control mice.** Daily profiles of food intake in 30-minutes intervals over 72 hours were determined by indirect calorimetry in 16-week-old Bmal1-iKO (**B**; n=5) and WT control (**A**; n=6) mice under ALF. The grey box indicates the dark phase.

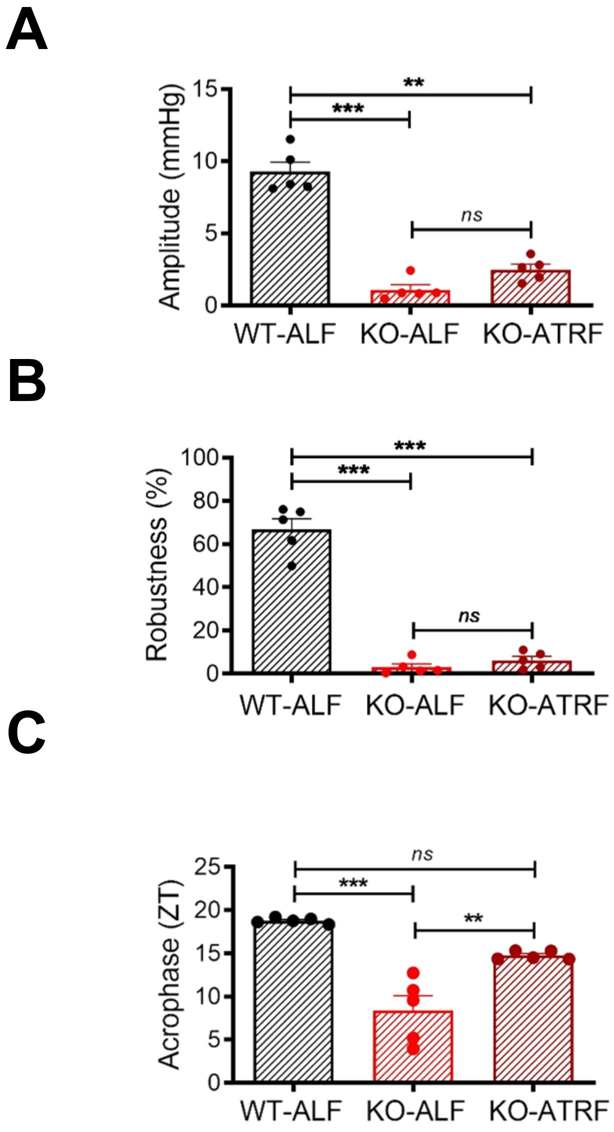


**Supplemental Figure 24. ATRF fails to restore mean arterial pressure (MAP) circadian rhythm in Bmal1-iKO mice.** Sixteen-week-old Bmal1-iKO and control (wild-type) mice were fed ALF or ATRF. MAP was measured by telemetry 2 days after ATRF. (**A**) Amplitude of the MAP circadian rhythm. (**B**) Robustness of the MAP circadian rhythm. (**C**) Acrophase of the MAP circadian rhythm. The data were analyzed by cosinor and 1-way ANOVA with multiple comparisons test. **, $p<0.01$; ***, $p<0.001$; ns, not significant.

**Supplemental Table 1. Real-time PCR primer information.**

| Gene | Primer | Sequence |
|---|---|---|
| Bmal1 | Forward | 5'-ATCAGCGACTTCATGTCTCC-3' |
| | Reverse | 5'-CTCCCTTGCATTCTTGATCC-3' |
| Clock | Forward | 5'-GGCGTTGTTGATTGGACTAGG-3 |
| | Reverse | 5'-GAATGGAGTCTCCAACACCCA-3' |
| Per1 | Forward | 5’-TCGAAACCAGGACACCTTCTCT-3’ |
| | Reverse | 5’-GGGCACCCCGAAACACA-3’ |
| Per2 | Forward | 5'-AGGCGTCCTTCTTACAGTGAA-3' |
| | Reverse | 5'-CAGGTTGAGGGCATTACCTCC-3' |
| Cry1 | Forward | 5'-TCGCCGGCTCTTCCAA-3' |
| | Reverse | 5'-TCAAGACACTGAAGCAAAAATCG-3' |
| Cry2 | Forward | 5’-CCTCGTCTGTGGGCATCAA-3’ |
| | Reverse | 5’-GCTTTCTTAAGCTTGTGTCCAGATC-3’ |
| Rev-erba | Forward | 5'-CCCTGGACTCCAATAACAACACA-3' |
| | Reverse | 5'-GCCATTGGAGCTGTCACTGTAG-3' |
| Rorc | Forward | 5’-TCCACTACGGGGTTATCACCT-3’ |
| | Reverse | 5’-AGTAGGCCACATTACACTGCT-3’ |
| Agt | Forward | 5’-TCTCTTTACCCCTGCCCTCT-3’ |
| | Reverse | 5’-CAGGCAGCTGAGAGAAACCT-3’ |
| Renin | Forward | 5’-TCAGGGAGAGTCAAAGGTTTCC-3’ |
| | Reverse | 5’-ACAGTGATTCCACCCACAGTCA-3’ |
| Ace | Forward | 5’-AGCCCAAGTGTTGTTGAACGA-3’ |
| | Reverse | 5’-TGGATACCTCCGTGCTTTTCT-3’ |
| Ace2 | Forward | 5’-TCCAGACTCCGATCATCAAGC-3’ |
| | Reverse | 5’-TGCTCATGGTGTTCAGAATTGT-3’ |
| At1a | Forward | 5’-CCAAGAAAGCCATCACCAGATC-3’ |
| | Reverse | 5’-TTTCTGGGTTGAGTTGGTCTCA-3’ |
| Adra1a | Forward | 5’-ATGAGGAGCCAGGATACGTG-3’ |
| | Reverse | 5’-TCTGACTTGTCGGTCTTGAGG-3’ |
| Adra1d | Forward | 5’-TCTCCGTAAGGCTGCTCAAG-3’ |
| | Reverse | 5’-AACCAGCACAGGACGAAGAC-3’ |
